# Wildfire smoke delays breeding and alters reproductive outcomes in cavity-nesting songbirds

**DOI:** 10.64898/2026.08.18.745532

**Authors:** Hayley A. Spina, Olivia V. Sanderfoot, Ravan Ahmadov, Robyn L. Bailey, Eric James, Alexandra Karambelas, Sierra Raby, Joe Siegrist, Andrew N. Stillman, Morgan W. Tingley

**Affiliations:** Department of Ecology and Evolutionary Biology, University of California, Los Angeles; Los Angeles, 90095, USA; Department of Integrative Biology, University of Guelph; Guelph, N1G 2W1, Canada; La Kretz Center for California Conservation Science, University of California, Los Angeles; Los Angeles, 90095, USA; Department of Ornithology, Natural History Museum of Los Angeles County; Los Angeles, 90007, USA; Cornell Lab of Ornithology; Ithaca, 14850, USA; NOAA Global Systems Lab; Boulder, 80305, USA; Northeast States for Coordinated Air Use Management; Boston, 02111, USA; Fielding School of Public Health, University of California, Los Angeles; Los Angeles, 90095, USA; Purple Martin Conservation Association; Erie, 16505, USA

## Abstract

Smoke from spring boreal wildfires increasingly impacts eastern North America, exposing breeding birds to hazardous air pollution that may impact reproductive outcomes. Using data collected in 2018–2025 from 70,979 monitored nests of four widespread cavity-nesting songbirds, we found strong evidence that smoke greatly delays egg laying and can extend incubation and nestling duration. We further found that while smoke is associated with increased clutch sizes, in some species smoke exposure strongly decreases hatching or fledging success. Our results demonstrate that extreme smoke can have wide-ranging impacts on breeding birds, from altering phenology to impacting fitness. While the exact mechanisms underlying these results remain elusive, the full suite of effects suggests that modifications to adult behavior under smoky conditions is the most likely cause. As fire regimes shift, birds and other wildlife are at greater risk of exposure to toxic smoke during the breeding season, which may further exacerbate the biodiversity crisis.

## Introduction

With wildfire regimes changing globally ^1^, wildfire smoke is increasingly recognized as an urgent human health crisis ^2^. Emerging research suggests that smoke pollution may also impact the health of non-human animals ^3–7^. Among wildlife species, birds are especially vulnerable to smoke due to their unique respiratory physiology. The avian lung-air sac system features unidirectional airflow, tightly intertwined air and blood capillaries, and cross-current gas exchange—adaptations that support highly efficient respiration but also compound the toxicity of air pollution for birds ^8^. Although industrial and urban air pollution have been linked to a wide range of adverse health outcomes in birds ^9^, few studies have considered the impacts of exposure to wildfire smoke ^4,6,7,10^ and how these effects may be contributing to the well-documented decline of avifauna ^7,11,12^.

In temperate North America, fires large enough to cause significant smoke pollution tend to ignite in densely forested ecosystems during the hottest and driest parts of the year, typically in the late summer to fall ^13^. Yet, as climate change causes wildfire seasons to expand earlier from historic norms ^14^, fires are increasingly likely to directly overlap with the seasonal reproduction of many organisms. In recent years, including 2026, record-breaking wildfires have repeatedly spread across the boreal forests in Canada, releasing extreme quantities of aerosol pollutants into the atmosphere (Fig. 1). These emissions severely degrade air quality in both Canada and the eastern United States due to long-distance transport of fine particulate matter (PM_2.5_, particles < 2.5 microns; ^13,15^), resulting almost immediately in measurable human health impacts ^16^. Given that North American wildfires historically occur outside the breeding season for most bird species ^13,17,18^, the recent spring-summer Canadian wildfires provide an opportunity to broadly investigate the impact of smoke exposure on avian breeding behavior and success—thus previewing the outcomes of an inevitably smokier future.

**Fig. 1:**
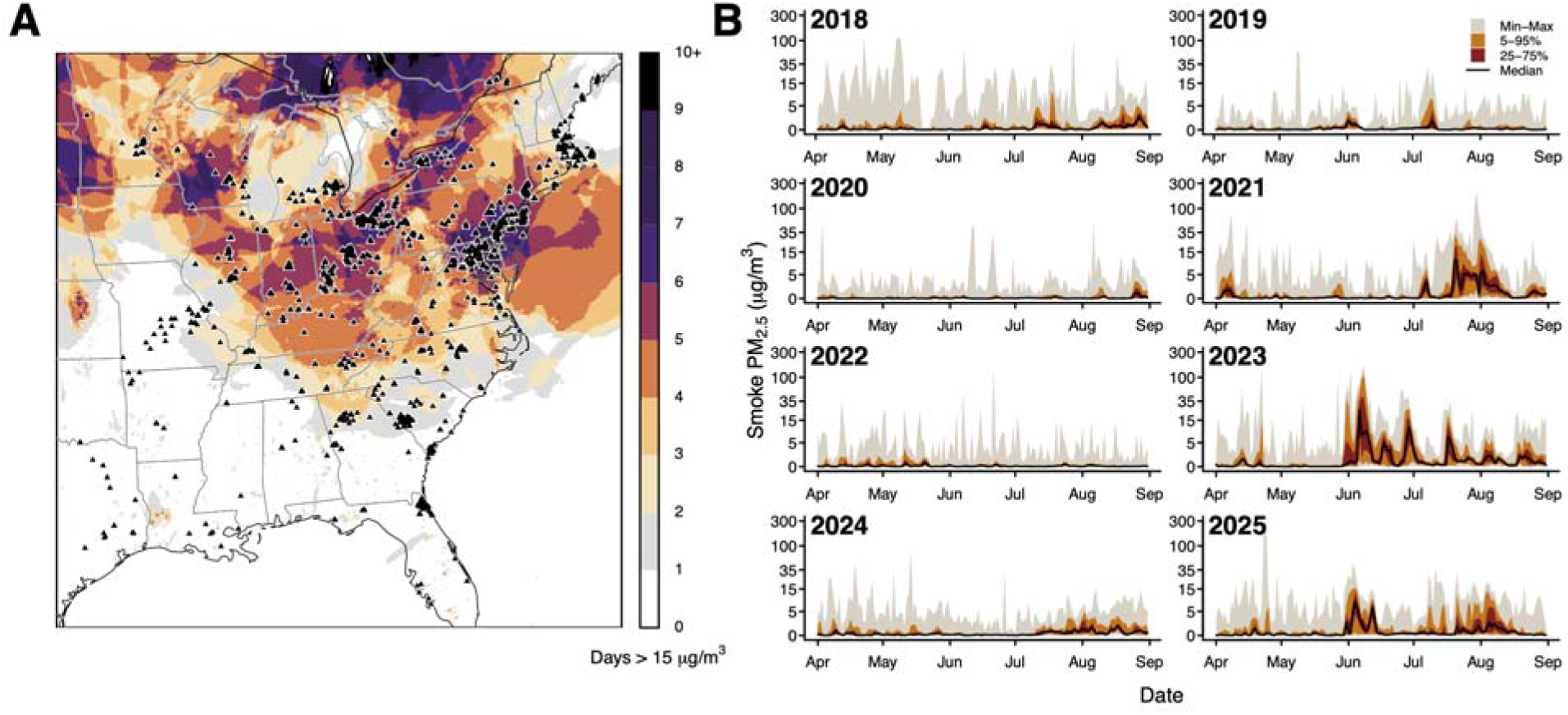
Smoke from boreal wildfires increasingly inundates eastern North America in spring-summer, resulting in widespread exposure to harmful air pollution coinciding with the avian breeding season. High-Resolution Rapid Refresh Smoke (HRRR-Smoke) data provide hourly estimates of fire-derived PM_2.5_ at a 3-km resolution. (A) The number of days in 2023 (01 April to 31 August) with smoke PM_2.5_ concentrations above 15 μg/m^3^—the World Health Organization’s recommended daily limit for humans—illustrates the spatial variation in smoke impacts for just one year with respect to the 6,505 monitored nest sites (triangles) for that year. See fig. S2 for all years. (B) While wildfire smoke during the North American avian breeding season is generally minimal, incursions during the breeding seasons of 2021, 2023, and 2025 repeatedly impacted air quality across the eastern U.S., sending daily smoke PM_2.5_ concentrations well above 15 μg/m^3^ for many of our monitored nest sites. Plots show daily quantiles of smoke PM_2.5_ concentrations experienced at all 9,619 nest monitoring locations, illustrating how smoke impacts on eastern temperate forests vary over time and space.

While limited observations suggest that wildfire smoke may disrupt bird breeding activity, the mechanisms are unclear ^7,19,20^. Anecdotal observations from the northeastern U.S. have supported hypotheses that smoke exposure may impair nestling growth ^21^ or affect breeding behavior ^22^, and a recent continental-scale study associates smoke with broad reductions in bird abundance in the following year ^7^. Studies have correlated other types of air pollution, such as emissions from industrial point sources, with reduced reproductive success in birds; for example, air pollution has been linked to reduced clutch size and hatching success in pied flycatchers (*Ficedula hypoleuca*; ^23^. Given that smoke contains many of the same air pollutants found in urban and industrial pollution and has already been linked to adverse health outcomes for birds ^4^, it is likely that wildfire smoke negatively impacts avian reproductive success. Yet, the extent to which smoke disturbance influences reproductive outcomes for birds is unknown.

Here, we leveraged two long-term North American nest-monitoring programs ^24,25^ to investigate how exposure to wildfire smoke affected avian breeding behavior and success in the eastern U.S. We compiled eight years (2018–2025) of data comprising 70,979 individual nesting attempts of four species—tree swallow (*Tachycineta bicolor),* purple martin (*Progne subis*), eastern bluebird (*Sialia sialis*), and northern house wren (*Troglodytes aedon*)—at 9,619 unique monitoring sites and merged these records with a state-of-the-art chemical transport model to evaluate nest-specific exposure to wildfire-associated PM_2.5_ (see Supplementary Materials; fig. S1 and table S1). We examined how smoke PM_2.5_ altered nest phenology and fitness at sites monitored repeatedly over time—both in years when smoke incursions heavily impacted air quality in the eastern U.S. (e.g., 2023; Fig. 1) and at the same locations in other years (figs. S2 and S3 and table S2). As such, our dataset takes advantage of both variations in smoke exposure over space (Fig. 1A and fig. S2) and within-sites over time (Fig. 1B and fig. S3). By considering exposure to smoke during the pre-laying, egg-laying, incubation, and nestling periods separately, we were able to test for effects of acute smoke on multiple phenological (i.e., first lay date, incubation duration, and nestling duration) and reproductive (i.e., clutch size, hatch success, and fledge success) outcomes (Table 1 and fig. S1), while controlling for spatiotemporal heterogeneity associated with specific nest sites or years (including climate and weather; see Supplemental Materials). We hypothesized that exposure to wildfire smoke would delay reproduction and lead to worse reproductive outcomes, including smaller clutches and reduced hatching and fledging success. Our results provide key insight into the wide-ranging impacts of wildfire smoke on avian reproduction and fitness and suggest that expected changes in the frequency, intensity, and seasonality of smoke disturbance may disrupt avian breeding behavior at wide-ranging scales.

**Table 1:**
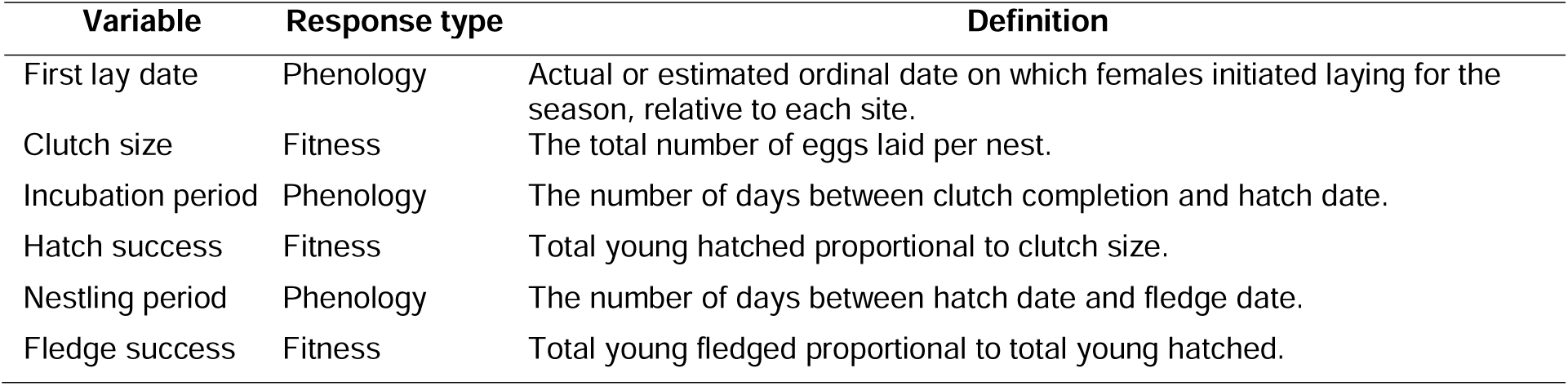
Definitions of the response variables analyzed to examine the ways in which wildfire smoke may influence the phenology and success of avian reproduction.

## Results

### Wildfire smoke delays egg laying and increases clutch size

We evaluated how exposure to wildfire smoke influenced the onset of egg-laying and found consistent evidence ^26^ of positive effects of smoke PM_2.5_ on lay date at nest sites (Fig. 2A and table S3). Specifically, we found very strong evidence that exposure to smoke PM_2.5_ during pre-laying (i.e., the fourteen days prior to the nest-specific average lay date) was associated with delayed lay dates for all species: eastern bluebird (sample size of nests (n) = 16548, p < 0.001, 95% confidence interval (CI) = [2.03, 2.88]), northern house wren (n = 4087, p < 0.001, 95% CI = [0.66, 2.68]), purple martin (n = 5313, p < 0.001, 95% CI = [0.58, 1.11]), and tree swallow (n = 18848, p < 0.001, 95% CI = [1.25, 1.59]). We also investigated the effect of smoke exposure during the laying period on clutch size (Fig. 3A and table S4). Contrary to our hypothesis, we found moderate to very strong evidence that exposure to smoke PM_2.5_ during egg laying increased clutch size for all species: eastern bluebird (n = 19946, p = 0.035, 95% CI = [0.001, 0.01]), northern house wren (n = 5262, p = 0.001, 95% CI = [0.004, 0.02]), purple martin (n = 25369, p < 0.001, 95% CI = [0.004, 0.01]), and tree swallow (n = 20387, p < 0.001, 95% CI = [0.004, 0.01]).

**Fig. 2:**
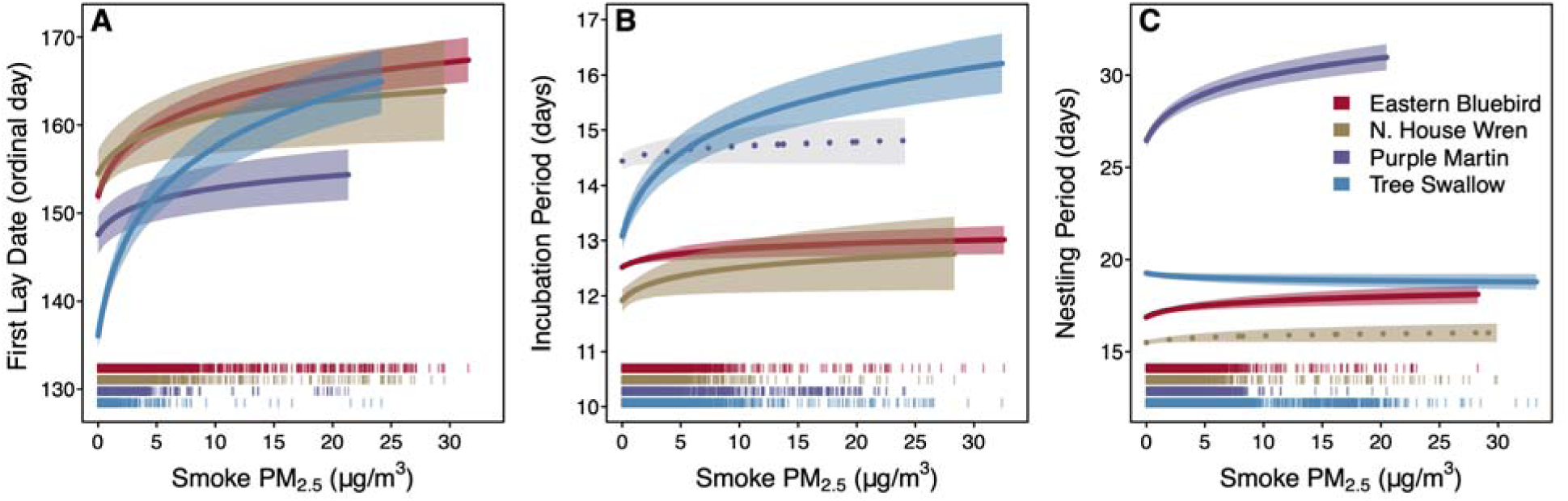
Smoke is associated with multiple phenological impacts to the avian reproductive cycle. Independent phenological responses include (**A**) first lay date (ordinal day of year), (**B**) duration of incubation period, and (**C**) duration of nestling period. Each response was tested independently across four species of cavity-nesting birds. In each plot, lines indicate mean responses and ribbons represent 95% confidence intervals of responses. Strength of evidence for modeled relationships is signified by a solid mean line (indicating p < 0.05; alternatively, dotted line indicates p > 0.05) and a colored ribbon (indicating p < 0.10; alternatively, gray ribbon indicates p > 0.10). Data rugs at bottom illustrate nest-specific exposure to smoke observed for each species.

**Fig. 3:**
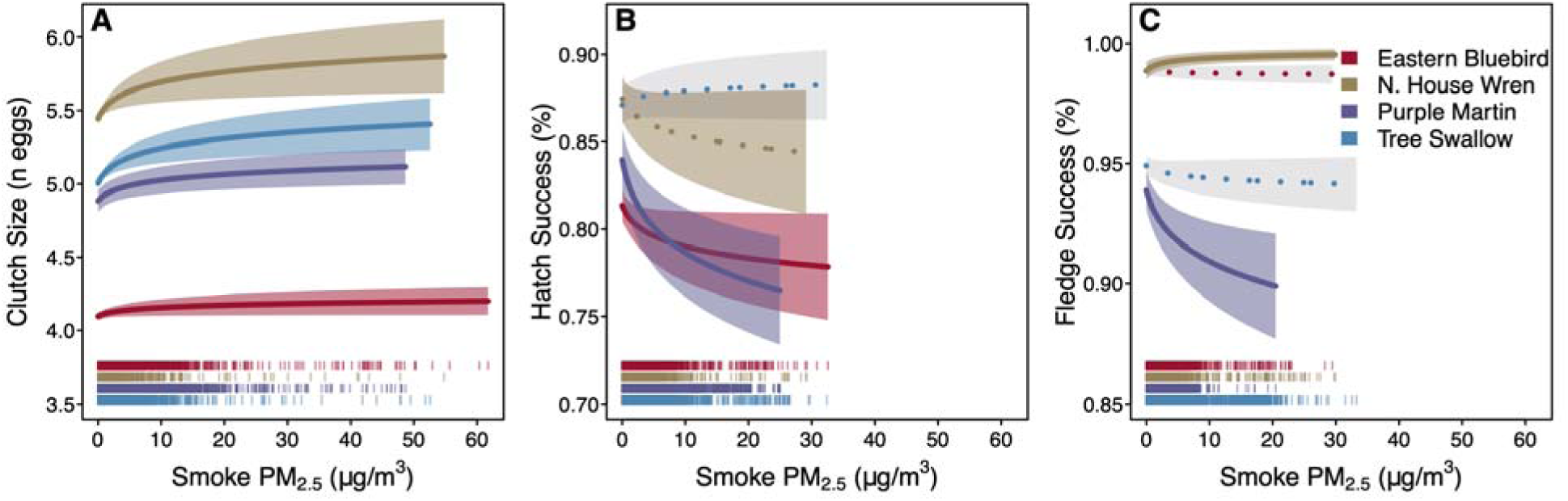
Smoke is associated with multiple fitness impacts to the avian reproductive cycle. Independent fitness responses include (**A**) clutch size, (**B**) hatch success, and (**C**) fledge success. Each response was tested independently across four species of cavity-nesting birds. Symbology as in Fig. 2.

### Wildfire smoke lengthens incubation and reduces hatch success

Exposure to smoke PM_2.5_ during incubation was strongly associated with delays in hatching for three species (Fig. 2B and table S5)—eastern bluebird (n = 11893, p < 0.001, 95% CI = [0.002, 0.01]), northern house wren (n = 3301, p = 0.017, 95% CI = [0.002, 0.02]), and tree swallow (n = 14190, p < 0.001, 95% CI = [0.02, 0.03])—but not purple martin (n = 4214, p = 0.114, 95% CI = [-0.001, 0.01]). We also tested the effect of smoke exposure during incubation on hatch success (Fig. 3B and table S6). We found very strong evidence of reduced hatch success for purple martin (n = 25333, p < 0.001, 95% CI = [−0.13, −0.07]), moderate evidence for eastern bluebird (n = 19095, p = 0.024, 95%, CI = [−0.06, −0.004]) and weak evidence for northern house wren (n = 4881, p = 0.077, 95% CI = [−0.09, 0.004]). We did not find evidence that smoke during incubation impacts hatch success of tree swallow (n = 19357, p = 0.258, 95% CI = [−0.01, 0.04]).

### Wildfire smoke delays fledging and may impact fledge success

We observed more variable effects of exposure to wildfire smoke during the nestling period. Exposure to smoke PM_2.5_ during the nestling period was very strongly associated with delays in fledging for two species (Fig. 2C and table S7)—eastern bluebird (n = 12996, p < 0.001, 95% CI = [0.006, 0.015]) and purple martin (n = 20602, p < 0.001, 95% CI = [0.02, 0.03])—and weakly associated with delays in fledging for northern house wren (n = 2922, p = 0.064, 95% CI = [0.001, 0.01]). In contrast, we found moderate evidence that smoke advances fledging for tree swallow (n = 12954, p = 0.049, 95% CI = [−0.01, −0.001]). We also tested the effect of smoke exposure during the nestling period on fledge success (Fig. 3C and table S8) and found very strong evidence of decreased fledge success for purple martin (n = 22201, p < 0.001, 95% CI = [−0.13, −0.05]) but increased fledge success for northern house wren (n = 3788, p < 0.001, 95% CI = [0.09, 0.25]). We found no evidence of fledge success effects for eastern bluebird (n = 14693, p = 0.440, 95% CI = [−0.06, 0.03]) or tree swallow (n = 15854, p = 0.178, 95% CI = [−0.07, 0.01]).

## Discussion

Our study reveals the consequences of breeding-season wildfire smoke events for four species of cavity-nesting songbirds in North America, providing large-scale evidence of pervasive effects of smoke pollution on avian fitness. We find that acute exposure to smoke is associated with delayed reproduction via postponement of egg laying and longer incubation periods.

Furthermore, we find evidence that acute smoke is associated with both increased clutch size and a general reduction in hatch success. Smoke appears to have more variable impacts on the length of the nestling period and fledge success, with strong evidence for negative effects on one species in particular, the purple martin. Altogether, our results suggest that wildfire smoke generally delays avian breeding behavior and can worsen reproductive outcomes for some cavity-nesting songbirds. Although this study is limited to four species, these findings provide mechanistic insight into recent suggestions that wildfire smoke may be contributing to ongoing large-scale population declines ^7,27^.

We find strong support for our hypotheses that wildfire smoke causes birds to postpone and lengthen reproduction. We found that smoke exposure during pre-laying, incubation, and nestling periods generally delayed egg laying, hatching, and fledging, respectively, to different degrees. For example, we estimate that eastern bluebirds exposed to extended periods of unhealthy air (i.e., average smoke PM_2.5_ concentrations above 15 ug/m^3^, the World Health Organization’s recommended daily limit for humans) delay laying eggs by 9.3 days (Fig. 2A), whereas exposure to smoke above the same threshold during incubation delays hatching by 0.3 days (Fig. 2B) and during nestling delays fledging by 0.8 days (Fig. 2C). Delayed first broods may reduce the probability of second clutch production in double-brooded species, such as eastern bluebirds and northern house wrens, resulting in negative consequences on annual fecundity and, potentially, lifetime reproductive success ^28–30^. Furthermore, deferred egg laying and lengthened incubation and nestling periods could lead to a phenological mismatch between the dietary needs of insectivorous birds and the seasonal abundance of their prey, resulting in reduced productivity ^31,32^. If so, delays in avian reproduction spurred by wildfire smoke could further desynchronize breeding activity from seasonal pulses in lower trophic levels, which are already shifting earlier at a faster rate than birds can match ^33,34^. However, wildfire smoke may also influence seasonal pulses in the availability of invertebrate prey; for example, exposure to smoke reduces survival and delays development of caterpillars, an important food resource for birds during the breeding season ^35^. As such, the delays in avian reproduction we observed could match shifts in insect availability. Although we find consistent evidence of delays in egg-laying and hatching, responses to smoke during the nestling stage were more variable – while eastern bluebirds and purple martins delayed fledging, tree swallows slightly advanced fledging. This suggests that not all species can or will respond to smoke disturbance in the same way across the life cycle.

Contrary to our expectations, we found consistent evidence that clutch size increases with exposure to smoke during the laying period. Although counter-intuitive, this intriguing finding has multiple non-exclusive explanations. First, initial delays in reproduction may have downstream effects that explain increased clutch size. Delayed egg laying has been shown to increase clutch sizes in multi-brooded species that breed in temperate regions ^36^, such as eastern bluebirds and northern house wrens. Multi-brooded species often initiate breeding in early spring, before conditions are optimal; thus, delays in egg laying can result in better alignment between hatching and peak resource availability for the first brood ^34,36,37^, which could lead to increased clutch size. Simultaneously, perceptions of smoke as a terminal risk by egg-laying females could shift cost-benefit trade-offs in reproductive investment, such that an uncertain present triggers increased reproductive investment in the current brood via future reproductive opportunities ^38^. Although this hypothesis has not previously been tested with smoke, multiple experiments support the theory that changes in terminal risk prompts increased clutch sizes in birds ^39,40^, including in northern house wrens ^41^.

Larger clutches may also partially explain our phenological results, in particular our finding that smoke increases the duration of incubation. Larger clutches with more eggs generally take longer to hatch, as incubation temperatures are more variable ^42,43^. However, eggs would also take longer to hatch if smoke exposure disrupted nest attendance and incubation behavior in adults ^44,45^. Additionally, smoke likely reduces ambient temperatures by blocking solar radiation ^46^, which may also delay hatching ^47,48^.

Critically, our results demonstrate that wildfire smoke can impact not just the timing of reproduction, but also its outcomes, including hatch success (Fig. 3B) and fledge success (Fig. 3C). For example, we estimate the probability of a purple martin egg hatching decreases by 4.5% when mean smoke exposure during incubation increases to 15 ug/m^3^, and the probability of an individual surviving to fledge decreases by 2.8% when exposed to the same smoke conditions during the nestling period. While negative impacts of smoke on reproductive success were not universally demonstrated in all study species—in particular, northern house wrens showed increased fledge success under smoky conditions (Fig. 3C)—the preponderance of evidence suggests that reproductive success of cavity-nesting birds declines with smoke exposure. There are several possible mechanisms to explain such fitness declines, including impacts on both parents and offspring. Generally, wildfire smoke may disrupt parental incubation behavior or decrease ambient temperature ^49^, both of which would decrease hatch success ^45^. Alternatively, although not well studied, toxic smoke particulates could directly impact offspring health, whether absorbed passively through eggshells or inhaled by nestlings ^6,50^.

The strength of evidence and general consistency in our findings across species suggest that birds are more vulnerable to smoke at earlier reproductive stages. For example, the effects of smoke on incubation duration are stronger and more consistent than on nestling duration; similarly, the effects on hatch success are stronger and more consistent than those on fledge success. This suggests that the strongest impacts are likely mediated by adult behavior, particularly the control that adults have over incubation, and further indicates that exposure to wildfire smoke may not be directly lethal to nestlings. However, sublethal impacts may be significant yet are not evaluated here—nestlings that survive to fledge may still be in poor condition ^21^ and could experience long-term health effects that influence lifelong fitness ^6^. Existing research suggests that a single wildfire smoke event experienced in youth can impair respiratory function in animals into adolescence and beyond ^51^. Future work should examine how wildfire smoke impacts the health of birds across life history stages.

While weather extremes such as heat waves, cold snaps, and rain events can influence the timing and success of avian reproduction—including having much stronger effects on fitness than those found here ^52,53^—the direction and magnitude of these effects vary by species, geographic location, and reproductive stage ^54,55^. Given that the most extreme smoke disturbances occurred during our study in June (Fig. 1), after the typical spring window for cold snaps and before extreme summer heat, it is unlikely that the smoke responses reported here are spatially and temporally coincident with independent climatic extremes. To evaluate this potential, however, we conducted a sensitivity analysis (see Supplementary Materials), finding the strongest correlations for smoke exposure with maximum April temperature ( = 0.05 to 0.12) and total May precipitation ( = −0.30 to −0.14; table S9). We then re-ran all six response models for each species with the inclusion of these two climatic variables and found that primary inference on all responses was unchanged (figs. S4 and S5 and tables S10 through S15), even after accounting for (often significant) impacts of climate.

Despite the general consistency in responses found across species, our study is limited to four species with similar life history traits; all four species feed on insects, nest in cavities, and undergo some seasonal migration. Future research may discover that birds with a different set of life history traits may vary in sensitivity to wildfire smoke. Some predatory birds have been known to use fire and smoke for hunting ^56^, whereas granivorous or frugivorous birds may experience fewer disruptions to foraging than insectivorous ones during smoke events.

Additionally, long-distance migrants may be more susceptible to negative smoke impacts than residents or short-distance migrants, as their breeding phenology is generally later and less flexible ^34^, potentially exposing them to more smoke during reproduction. Determining how various reproductive behaviors and life histories align with expected wildfire smoke exposure under current and emerging fire and smoke regimes is a key next step in evaluating species-specific ecological ^6^ and evolutionary ^57^ consequences of wildfire smoke.

In summary, wildfire smoke has emerged as a defining public health issue in the 21^st^ century ^2^, yet few studies have considered how a smokier future will impact animal populations. Although some of the effect sizes we report are small, impacts are being experienced across an enormous scale—nearly the entire breeding ranges of our focal species. Specifically, the 2023 breeding season (April–June) witnessed particularly high smoke exposure across eastern temperate forests, overlapping with the abundance distributions of 98% of northern house wrens, 92% of tree swallows, 57% of eastern bluebirds, and 32% of purple martins present in our study region (see Supplementary Materials). Using established population size estimates ^58^, we estimate that across the four focal species, more than 25.1 million individuals (95% CI: [23.4 million, 27.6 million]) in our study region were exposed to smoke concentrations greater than 15 μg/m^3^ in June 2023 alone. In this context, even one anomalous wildfire smoke event has the potential to disrupt avian reproduction at a semi-continental scale.

## Conclusion

While ecologists have long understood the transformative impacts of wildfire on ecosystems, wildfire smoke is only beginning to be understood as an ecological disturbance in and of itself, with impacts that differ markedly in scale and strength ^6^. Although our results demonstrate how wildfire smoke can delay avian breeding phenology and even impact reproductive success, the specific mechanisms underlying these impacts are unclear. Given ongoing and alarming declines in bird abundance across North America ^11,12^, our findings emphasize the urgent need to ramp up research on the effects of wildfire smoke on avian and wildlife health. We also note that boreal and temperate conifer fire seasons are rapidly becoming more extreme ^59^, suggesting that breeding season smoke events are likely the new normal. Given the unpredictable nature of smoke disturbance, large-scale and long-term monitoring and participatory science initiatives will be crucial to our future understanding of wildfire smoke impacts. Such future studies will also benefit from interdisciplinary collaborations between ecologists and atmospheric scientists to develop smoke exposure data products specifically for wildlife health applications.

## Methods

### Breeding Bird Observations

Our study focused on observations of breeding birds in the temperate forests of the eastern U.S. Eastern temperate forests were variably impacted by smoke from Canadian wildfires between May and July in the years 2018–2025 (Fig. 1 and fig. S2).

We obtained records of breeding birds from two long-term, participatory science programs – NestWatch and Project MartinWatch. NestWatch (www.nestwatch.org; ^24^) has been administered by the Cornell Lab of Ornithology since 2008 to engage the public in collecting standardized data on the reproductive activity of birds. The open-access dataset contained > 860,000 nest records across all species at the time of access. While NestWatch accepts data from any location globally, 96% of records are from the U.S. ^24^. Project MartinWatch (www.purplemartin.org) has been run by the Purple Martin Conservation Association since its inception in 1995 to monitor the reproductive activity of purple martin in provisioned housing ^60^. This ongoing, longitudinal study has tracked the fate of more than 140,000 nests of this one species. These data represent hundreds of thousands of hours of effort from thousands of participatory scientists. Both programs collect standardized data on species, location, clutch size, egg-laying date, hatch date, fledge date, and outcome for each monitored nest.

While these programs each execute a series of validation steps to improve data accuracy ^24^, additional checks are recommended to reduce potential errors (https://nestwatch.org/explore/nestwatch-open-dataset-downloads/). As such, we implemented a series of quality control steps recommended by Taff and Shipley ^55^ as part of our workflow.

First, we manipulated the NestWatch and Project MartinWatch datasets into a common format to enable the merging of records. Then, we filtered our dataset to include only nests located within the eastern temperate forest ecoregion. To do so, we downloaded shapefiles representing level I ecoregions of North America from the U.S. Environmental Protection Agency ^61^. We then used the *sf* package ^62^ in R ^63^ to match nests to ecoregions based on their GPS locations (i.e., latitude and longitude coordinates) and filtered the dataset to include only nests observed within the eastern temperate forests of the U.S. between 2018 and 2025.

Our analysis required that all nest records include a lay date (Table 1). For records missing lay date, we estimated lay date using other reliable information when provided. Following Taff and Shipley ^55^, we inferred approximate lay dates for nests for which eggs hatched by subtracting the reported clutch size (i.e., number of eggs) and the average incubation length (i.e., number of days) from the hatch date, since most birds lay one egg per day prior to incubation. Average incubation lengths were extracted from species accounts included in *Birds of the World* ^17^. We were unable to estimate lay date for nests also missing a valid hatch date and clutch size (“invalid” records include clutch sizes that were blank, negative, or zero); as such, we removed these records from the dataset. After ensuring that all records included a lay date (observed or estimated), we temporally filtered the dataset to include nests with lay dates between May 1 and July 31 (to align with a smoke exposure period of 15 April to 31 August). Next, we removed records of nests that failed due to invasive species management (e.g., participants removed nests of House Sparrows, *Passer domesticus*, or European Starling, *Sturnus vulgaris*, in nest boxes) and records for which the reported year did not match that of the lay date (indicative of data collection errors in our study region).

After implementing these universal filtering steps, we chose our study species based on data availability. We focused our analysis on species with at least 200 nest records in each of our study years, 2018–2025. Since smoke from boreal wildfires in Canada variably degraded air quality across our study area at different times in different years (Fig. 1 and fig. S2), this step ensured that we had sufficient data to consider how reproductive success varied across a wide range of smoke concentrations and locations.

An important note about these datasets is that individually monitored nests (often artificial nestboxes) are grouped within monitoring locations (’sites’). Each site has a unique spatial location. For NestWatch data, each unique nest location is treated as an individual ‘site’, meaning data contributors register each nestbox separately. For Project MartinWatch, the unique location is tied to the data contributor who registers a street address associated with the data.

Since purple martin almost exclusively nest colonially in multi-room ‘houses’, a single Project MartinWatch site will contain multiple nest records per year which generally represent separate nesting attempts by different pairs. In all cases, the data used here are distinguished by site and nest, not individual pair, so we cannot distinguish conclusively between double-brooding or multiple pairs. As data are gathered on individual nesting attempts and aggregated at the site level, our analysis uses the nest as the unit of inference and treats multiple nests per site within the same year as pseudoreplicates.

Our final universal filtering step was to include only nest records from sites monitored for at least three (consecutive or non-consecutive) years between 2018 and 2025 (table S1). While this resulted in discarding data from many sites, it ensured that our primary inference would focus on interannual changes in nest outcomes at repeated, individual sites as a function of wildfire smoke, rather than potential spatial signals. Our final dataset included nest records of four species: eastern bluebird (*Sialia sialis*; n = 19,953 nests at 4,115 sites), northern house wren (*Troglodytes aedon*; n = 5,267 nests at 1,074 sites), purple martin (*Progne subis*; n = 25,371 total nests across all years at 1,018 sites), and tree swallow (*Tachycineta bicolor*; n = 20,388 nests at 4,294 sites). We used this final filtered dataset to generate six subsets of nest records for our analysis, each associated with a breeding variable of interest (fig. S1 and Table 1), including (1) first lay date, (2) clutch size, (3) incubation period, (4) hatch success, (5) nestling period, and (6) fledge success. For each subset, we implemented additional filtering criteria and/or data processing steps as described below:

#### First Lay Date

Within our study period (May 1 – July 31), we retained only the first nest records for each species at a given site within a given year.

#### Clutch Size

We removed nest records with unrealistic clutch sizes (i.e., larger than the typical range for each species) using the cut-offs suggested by Taff and Shipley ^55^.

#### Incubation Period

First, we removed nest records for which the reported year did not match that of the hatch date and records for which the hatch date reported was earlier than the estimated incubation start date (indicative of data collection errors). Second, we removed nest records that were missing a hatch date. Then we removed records for which no young hatched. Finally, we removed nest records with an incubation duration outside the 1^st^ and 99^th^ species-specific quantile within our dataset (indicative of data collection errors).

#### Hatch Success

We first implemented the initial filtering step described above for incubation period. Next, we removed records with an unrealistic number of young to hatch (e.g., larger than the typical range for each species), using the cut-offs suggested by Taff and Shipley ^55^. We also filtered out records of nests missing information (i.e., blank or NA) on the number of young that hatched. Finally, we removed records for which the reported number of young hatched was negative or greater than the documented clutch size (indicative of data collection errors).

#### Nestling Period

We first considered nest records where eggs hatched but which were missing hatch date. For these records, we estimated hatch dates by adding the clutch size (i.e., number of eggs) and average incubation length (i.e., number of days) to the lay date ^55^. We then, as with incubation period, checked for year mismatches between lay date, hatch date, and fledge date, and additionally removed nests for which these three reported dates were out of order (indicative of data collection errors). We also removed nest records with a nestling duration outside the 1^st^ and 99^th^ quantile within our dataset (indicative of data collection errors). Finally, we filtered out records of nests for which no young fledged and records which were missing a fledge date.

#### Fledge Success

We first considered records for which information on fledge date was missing but where successful fledging was reported. In such cases, we computed the average length of the nestling period for each species using our dataset (*Progne subis* = 28 days; *Tachycineta bicolor* = 19 days; *Sialia sialis* = 17 days; *Troglodytes aedon* = 15 days), then estimated missing fledge dates by adding the average nestling period to the hatch date. We then implemented all but the last two filtering steps described above for the nestling period variable. We then removed records with an unrealistic number of fledglings (e.g., larger than the typical range for each species), using the cut-offs suggested by Taff and Shipley ^55^. We also filtered out records of nests missing information (i.e., blank or NA) on the number of young that fledged. Finally, we removed records for which the reported number of young fledged was negative or greater than the documented clutch size or number of young hatched (indicative of data collection errors).

As a final filtering step applied to all data subsets, we additionally checked and removed any sites with fewer than three years with nesting data. Due to differing filtering steps, each response variable subset comprised different sample sizes of nests at sites in each year (table S2). Data for first lay date included the most sites, but clutch size included the most unique nests (table S2).

Responses that required greater filtering and data quality requirements generally retained fewer nests at fewer sites, but always a minimum of 200 nests per year per species per response (table S2). Filtered and cleaned subsets of raw data used in our analysis are provided on Dryad (http://datadryad.org/share/LINK_NOT_FOR_PUBLICATION/rJqJeq613xpzAvpqhTn2i-LN3LFqauM2zrZ7gXvtya4).

#### Smoke Exposure

To estimate exposure to smoke at nest site locations, we used forecasts from the High-Resolution Rapid Refresh Smoke (HRRR-Smoke) model, developed by the National Oceanic and Atmospheric Administration (NOAA) Global Systems Laboratory ^64–66^. HRRR-Smoke simulates PM_2.5_ emitted by biomass burning by using satellite detections of fire radiative power, providing estimates of smoke-specific PM_2.5_ at a 3-km resolution across the continental U.S. ^65,66^.

Importantly, HRRR-Smoke also accounts for smoke transported from Canada and other surrounding land areas by taking boundary conditions from the 13-km Rapid Refresh (RAP) model, which covers all of North America ^65,67^. We extracted hourly modeled near-surface smoke PM_2.5_ and averaged the hourly forecasts to calculate daily mean concentrations of smoke-specific PM_2.5_ across the lower 48 U.S. states ^68,69^. Some daily averages could not be created due to missing model outputs, particularly during the years HRRR-Smoke was considered experimental (2018 and 2019). The count of missing days per month in our dataset ranges from 0 to 5 (mean = 0.725). Next, we extracted daily average smoke PM_2.5_ concentrations at each nest location for each day from April–August over the years 2018–2025. We excluded 2 days (5 and 6 June, 2019) for which HRRR-Smoke predicted concentrations exceeded 500 ug m^-3^, which were understood to be model-based outliers. To determine whether smoke-specific PM_2.5_ affected avian breeding success, we created exposure variables representing average smoke PM_2.5_ concentrations during four specific reproductive stages: pre-laying, laying, incubation, and nestling (figs. S1 and S3). To reduce potential circularity between first lay date and smoke exposure during the pre-laying period, we defined the pre-laying period as 14 days prior to the multi-year average first lay date for each site. For all other reproductive stages (i.e., laying, incubation, and nestling), we calculated exposure windows based on year-specific dates for each nest (i.e., late date, hatch date, fledge date; reported or estimated as described above).

The units of the smoke field in RAP, which covers North America with 13-km grid spacing and provides boundary conditions to HRRR-Smoke, were changed from ug m^-3^ to kg m^-3^ in late 2021; however, due to an oversight, the HRRR-smoke boundary conditions continued to be treated as ug m^-3^, leading to smoke boundary conditions a factor of 10^9^ too small. This issue was corrected in the operational HRRR-Smoke on 6 June 2023. Since 2022 was a relatively inactive fire year in Canada, the problem is unlikely to significantly impact the 2022 results. However, HRRR-Smoke likely underestimated smoke concentrations in May 2023 when there were major fires occurring in portions of Canada outside the HRRR domain. Thus, the 2023 HRRR-smoke PM_2.5_ estimates should be considered conservative prior to 6 June of that year.

The HRRR-Smoke model is uniquely valuable to public health and ecological smoke studies due to its specificity to exposure-relevant near-surface PM_2.5_ as well as its exclusion of industrial emission and other non-smoke PM_2.5_ contributions. Evaluation during the 2018 Camp Fire in California found that HRRR-Smoke captured the magnitude and spatial area of the wildfire smoke compared to surface observations, although it underpredicted smoke during the later portion of the event when fire radiative power was underestimated by satellites due to clouds and thick smoke ^70^.

#### Statistical Analysis

For each species, we built six generalized linear mixed-effects models to test for effects of smoke PM_2.5_ on (1) first lay date, (2) incubation period, (3) nestling period, (4) clutch size, (5) hatch success, and (6) fledge success, while controlling for site- and year-specific effects. We modeled first lay date as the ordinal day of year with a Gaussian distribution and identity link function.

Incubation period, nestling period, and clutch size were modeled with a zero-truncated Poisson distribution and log link. We used binomial models with a logit link to analyze hatch success and fledge success (i.e., accounting for the variable denominator of each nest record). All models were built using the *glmmTMB* package ^71^ in R ^63^.

For each response variable, we examined the impact of the average smoke PM_2.5_ exposure during a specific time window relevant to the particular response (fig. S1). For first lay date we used the site average pre-laying period (range: <0.001–31.57 µg/m³); for clutch size we used the laying period (range: <0.001–61.73 µg/m³); for incubation duration and hatch success we used the incubation period (range: <0.001–32.56 µg/m³); and for nestling duration and fledging success we used the nestling period (range: <0.001–33.25 µg/m³). To minimize the leverage of extreme smoke values, to account for hypothesized non-linear impacts, and to account for zero values, we log(x + 1) transformed PM_2.5_ exposure variables. We then took each transformed PM_2.5_ exposure variable and centered and scaled it to a standard deviation of 1, to aid model fitting. To account for expected impacts of seasonality on reproductive success ^34^, we included the ordinal day-of-year as orthogonal linear and quadratic terms (with the *poly* function in ^63^) in all models except for first lay date. For models of clutch size, incubation period, hatch success, nestling period, and fledge success, we used first lay date, incubation start date, hatch date, hatch date, and fledge date, respectively, as the reference day of year.

We included a random effect of year to account for interannual variability in breeding success and a random effect of site (i.e., location ID) to account for differences in local habitat, weather, geographical location, and/or resource availability across sites that may have influenced reproductive outcomes. Random effects also help account for potential pseudoreplication (i.e., multiple nests at the same site in a given year), and the non-independence of nests monitored at the same site over multiple years. Validation of all models was assessed using the *DHARMa* package ^72^. Complete model results for each response and species are provided in Tables S3–S8.

Model-informed inference focuses on both presentation of effect sizes and evaluation of strength of evidence as indicated by *p*-values. Rather than use a single, arbitrary cut-off of “significance,” we employ the language of ‘evidence’ (following ^26^), using the terms ‘weak,’ ‘moderate,’ ‘strong,’ and ‘very strong’ to describe *p*-values falling below 0.1, 0.05, 0.01, and 0.001, respectively ^26^.

#### Sensitivity Analysis to Alternative Climatic Factors

Reproductive success in passerine birds can vary from year to year due to a multitude of factors, including environmental variation (e.g., rainfall, temperature). This effect has been shown with many of the species included in our analysis, even with the same datasets as used here ^54,55^. In part to account for climatic effects, all of our models for the main analysis include random intercepts for year (which controls for annual climatic patterns common across the eastern temperate forest study area) and site (which controls for climatic and microclimatic factors unique to each monitored location). However, these random effects do not account for interannual variation in weather and climate unique to each site. Nevertheless, since heavy smoke itself can impact local weather (e.g., via raising albedo which causes local cooling ^49^— which would be part of the total smoke impact we wished to measure), it was inappropriate to include weather variables in our models exactly matched to the reproductive periods during which smoke was measured (see *Smoke Exposure*). Thus, we employed a sensitivity analysis to examine the sensitivity of our results to the inclusion of climatic factors.

We downloaded monthly gridded climate data from TerraClimate v1.1 (https://www.climatologylab.org/terraclimate.html; ^73^), which presents downscaled and interpolated time-varying data from ERA5 at a high spatial resolution (∼4km) and good accuracy. We focused on three variables (maximum temperature [Tmax], averaged monthly; minimum temperature [Tmin], averaged monthly; precipitation [Precip]; monthly total) and the months of April and May, as avian phenology and reproduction is primarily sensitive to early-season climate ^34,55^.

As a first pass analysis, we calculated the Pearson correlation between the smoke exposure for each of our six reproductive variables and the spatially- and annually-matched climate for each of the four climate variables. For this first analysis, we combined all species together. Across all variable combinations, correlation coefficients were low (|| 0.35; table S9), ranging from a high of 0.124 (Tmax-April with first lay date) to a low of −0.300 (Precip-May with fledge success).

To additionally examine the sensitivity of our smoke impact estimates to the inclusion of climatic variables, we re-ran all 24 GLMM models (see *Statistical Analysis*) with the additional inclusion of two covariates—Tmax-April and Precip-May—matched to the location and year of every monitored nest. All other analytical aspects of models remained the same. The results of these models (figs. S4–S5 and tables S10–S15) were largely equivalent to those without climate variables (Figs. 3–4 and tables S3–S8). In several cases, the strength of evidence changed: (i) purple martin shows no support for a relationship of smoke with incubation period duration in a model without climate (Fig. 2B) but moderate evidence for increased duration with climate (fig. S4B); (ii) northern house wren shows weak support for an increase in nestling period duration with smoke in a model without climate (Fig. 2C), but moderate support when including climate (fig. S4C); (iii) tree swallow shows moderate support for a decline in nestling period duration with smoke in a model without climate (Fig. 2C), but no support when including climate (fig. S4C); and (iv) northern house wren shows weak support for a decrease in hatch success with smoke in a model without climate (Fig. 3B), but no support for a relationship when including climate (fig. S5B). Inference on all other parameters and smoke relationships remained unchanged.

#### Quantifying Smoke Exposure Across the Eastern Temperate Forest

To assess how the effects of wildfire smoke on avian nesting ecology may scale across the eastern temperate forest ecoregion, we calculated the percent of each focal species’ population exposed to smoke concentrations > 15 µg/m³ during June 2023, a time of particularly widespread and intense smoke incursions to the eastern U.S. (Fig. 1). First, we summarized daily smoke concentrations from HRRR–Smoke into a raster representing the maximum concentration reached by each pixel in June 2023. We then applied a threshold of > 15 µg/m³, the World Health Organization’s guideline, to delineate the portion of the ecoregion exceeding this threshold as a single polygon. Next, we used relative abundance data products from eBird Status and Trends ^74^ to calculate the percent of each species’ population within the study region that overlapped the smoke polygon. We accessed relative abundance predictions via the R package *ebirdst* ^75^ and used the breeding season, 3×3-km data product for each focal species. To calculate exposure, we divided the sum of relative abundance within the smoke-affected area by the sum of relative abundance within the eastern temperate forest study region. The resulting number is the percent of each species’ eastern temperate forest population exposed to smoke concentrations > 15 µg/m³ during June 2023.

To calculate the total number of individuals exposed to smoke in our study region, we used regional adult population estimates obtained from the Partners in Flight Population Estimates Database ^58^ and as described and validated in ^11,76^. First, we calculated the proportion of each species’ U.S. and Canada population within the June 2023 smoke polygon using eBird relative abundance ^74^. Second, we multiplied this proportion by the U.S. and Canada population size estimate and associated upper and lower confidence intervals. The resulting number represents the estimated total number of adults (of any sex) exposed to smoke concentrations > 15 µg/m³ during June 2023.

## Acknowledgments

We are grateful to the many participatory scientists who contribute to NestWatch and Project MartinWatch and to the Cornell Lab of Ornithology and the Purple Martin Conservation Association for developing, funding, and managing these programs, respectively. Previous versions of the manuscript benefitted from review by the Tingley Lab as well as three anonymous reviewers. The scientific results and conclusions, as well as any views or opinions expressed herein, are those of the authors and do not necessarily reflect those of NOAA or the Department of Commerce.

## Author contributions

Conceptualization: HAS, OVS, MWT

Methodology: HAS, OVS, EJ, RA, SR, AK, ANS, MWT

Investigation: HAS, OVS, ANS, MWT Visualization: HAS, OVS, EJ, MWT Project administration: RLB, JS Supervision: MWT

Writing – original draft: HAS, OVS, MWT

Writing – review & editing: HAS, OVS, EJ, RA, SR, AK, RLB, JS, ANS, MWT

## Data and Code Availability

Raw data from NestWatch is publicly available at https://data.mendeley.com/datasets/wjf794z7gc/6. Raw data from Project MartinWatch is available upon request from JS. All filtered and processed datasets used in this study are available on Dryad at:

## Supplementary Materials for

### Supplementary Materials include

http://datadryad.org/share/LINK_NOT_FOR_PUBLICATION/rJqJeq613xpzAvpqhTn2i-LN3LFqauM2zrZ7gXvtya4

---

**Fig. S1.**
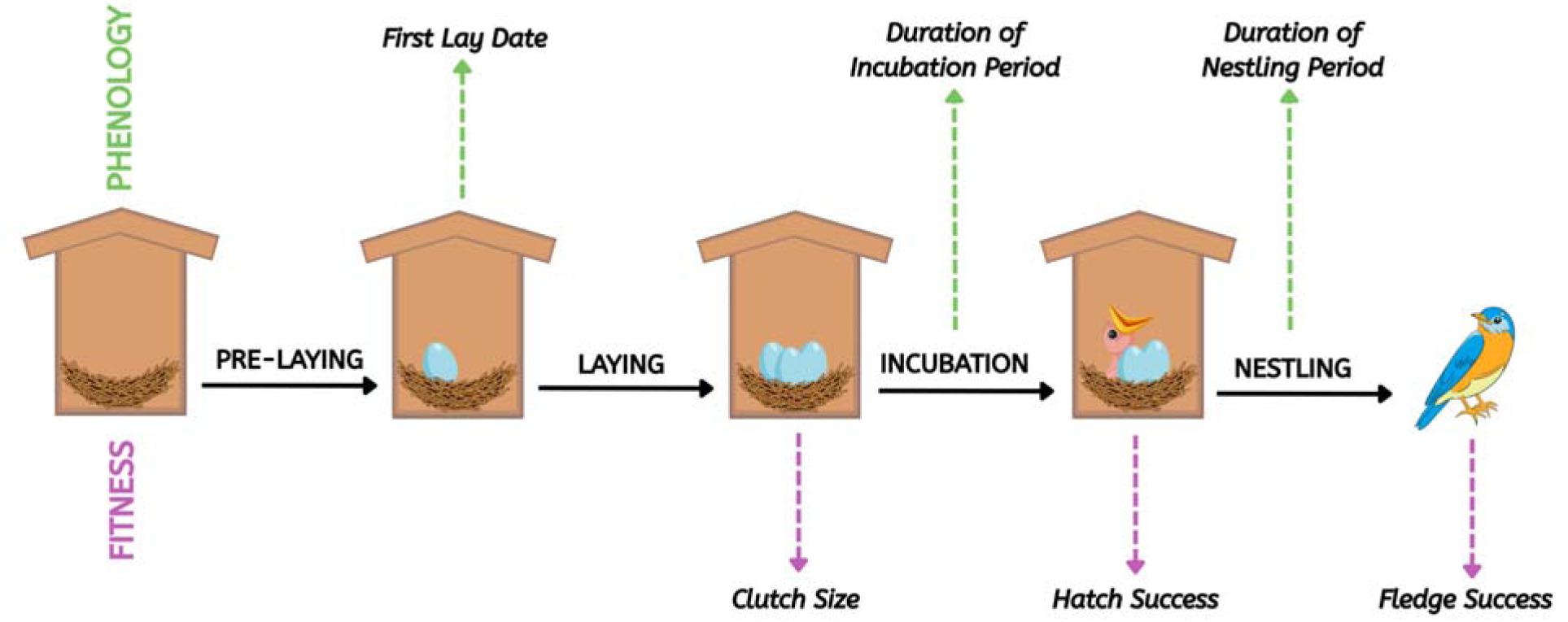
Standardized North American nest monitoring schemes provide opportunities to test both phenological and fitness impacts of wildfire smoke on the avian life cycle. Phenological impacts include first lay date, duration of incubation period, and duration of nestling period; fitness impacts include clutch size, hatch success, and fledge success. Each response variable is evaluated relative to average daily wildfire smoke occurring at the nest location during the reproductive interval immediately preceding the response (i.e., pre-laying, laying, incubation, or nestling periods).

**Fig. S2.**
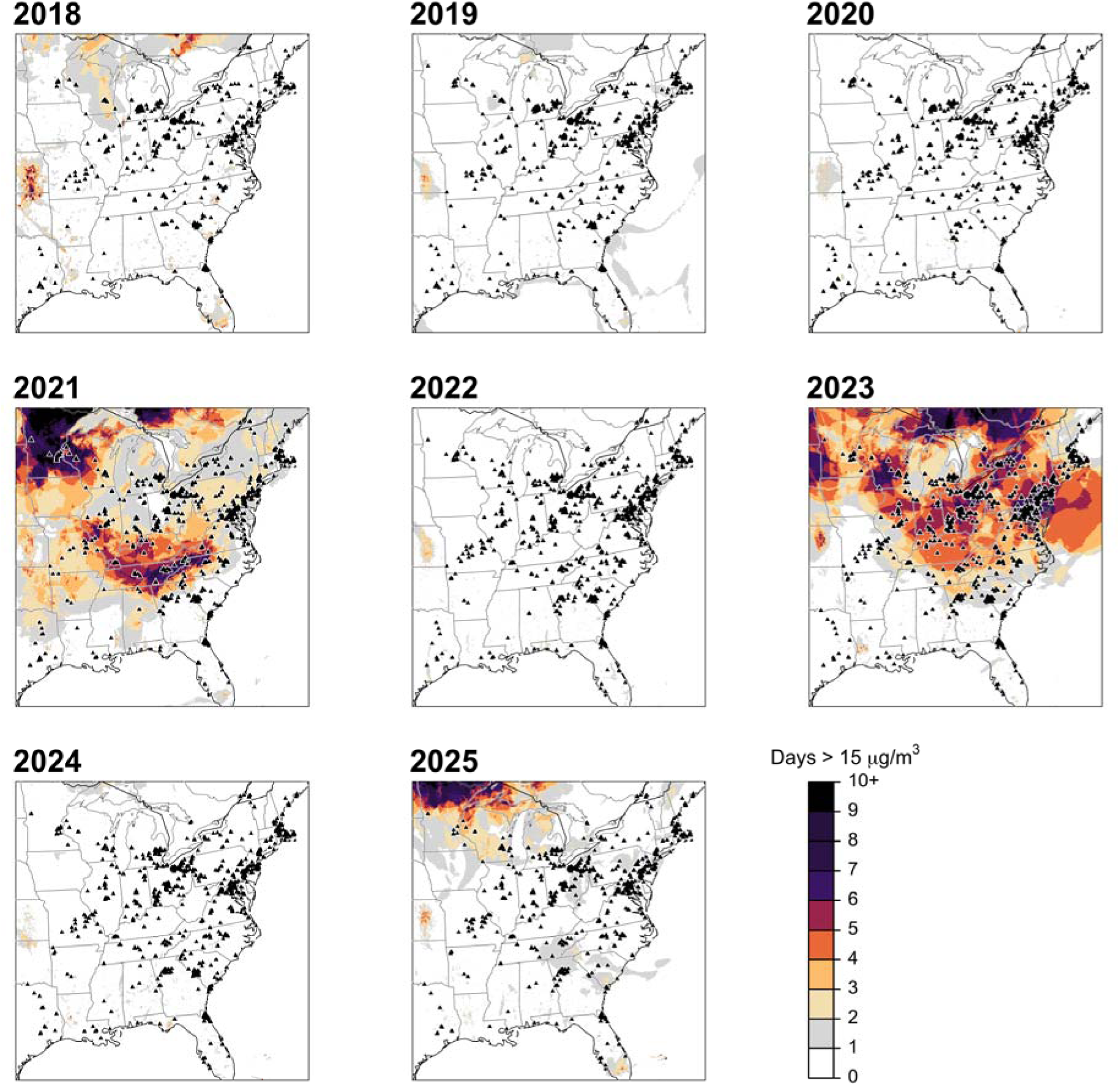
High-Resolution Rapid Refresh Smoke (HRRR-Smoke) data demonstrate the spatial variability in smoke impacts from year to year. Each annual panel shows the number of days each year (01 April to 31 August) with smoke PM_2.5_ concentrations above 15 μg/m^3^—the World Health Organization’s recommended daily limit for humans. Overlaid in each panel are the locations of monitored nest sites (all species combined; triangles) used for analysis in each year.

**Fig. S3.**
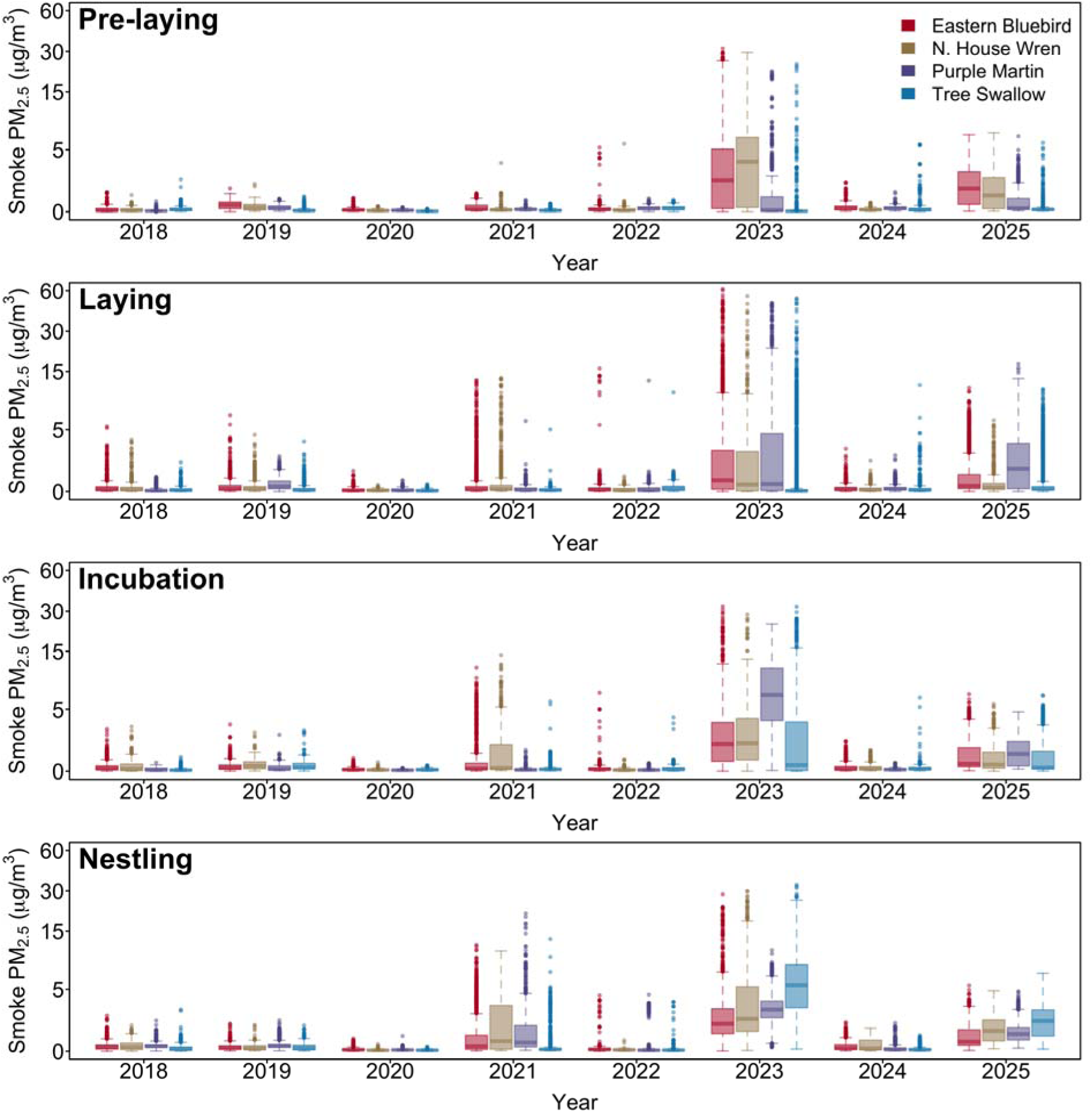
Smoke exposure during the Nearctic avian breeding season, 2018–2025. Boxplots (and points outside 1.5x interquartile-range) show HRRR-Smoke values of fire-attributed PM_2.5_ at monitored nest sites for all four species during each of the four reproduction periods (see fig. S1). Smoke was particularly present during avian reproduction in 2021, 2023, and 2025 (fig. S2), with widespread anomalously-high concentrations of smoke PM_2.5_ observed in eastern temperate forests of North America during the 2023 Canadian wildfires.

**Fig. S4:**
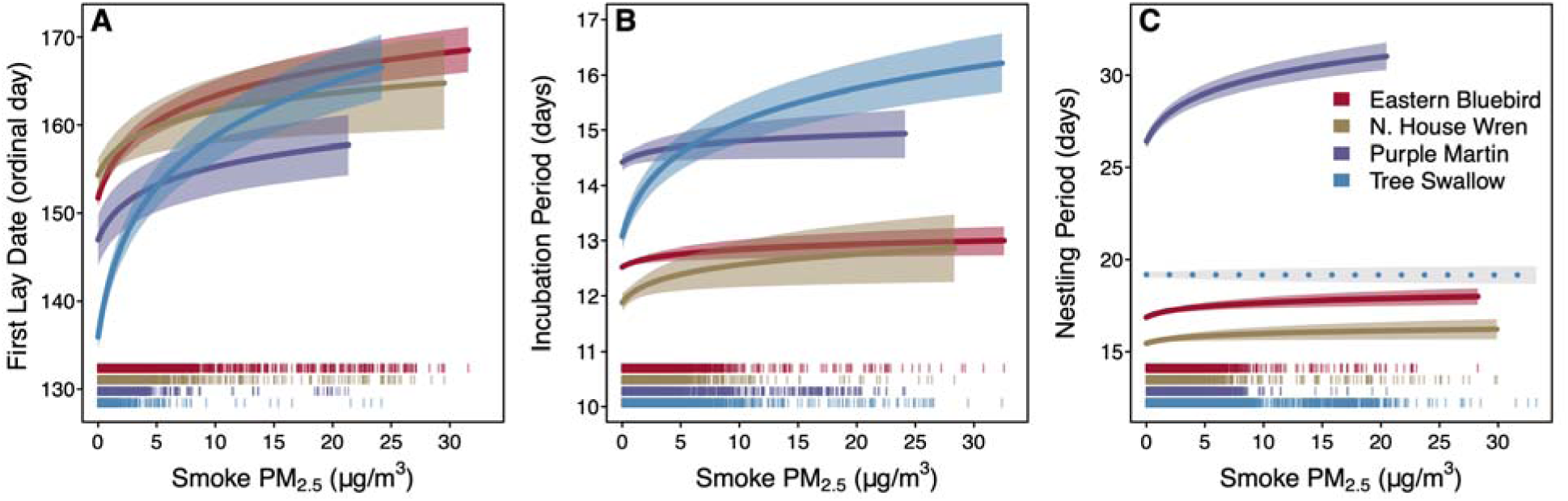
Smoke, controlling for site-specific spring climate, is associated with multiple phenological impacts to the avian reproductive cycle. Independent phenological responses include (**A**) first lay date (ordinal day of year), (**B**) duration of incubation period, and (**C**) duration of nestling period. Each response was tested independently across four species of cavity-nesting birds while controlling for maximum April temperature and total May precipitation at each site in each year (all responses), as well as the linear and quadratic phenological impact of day of year (B and C, only). In each plot, lines indicate mean responses and ribbons represent 95% confidence predictions of responses. Strength of evidence for modeled relationships is signified by a solid mean line (indicating p < 0.05; alternatively, dashed line indicates p > 0.05) and a colored ribbon (indicating p < 0.10; alternatively, gray ribbon indicates p > 0.10). Data rugs at bottom illustrate nest-specific exposure to smoke observed for each species.

**Fig. S5:**
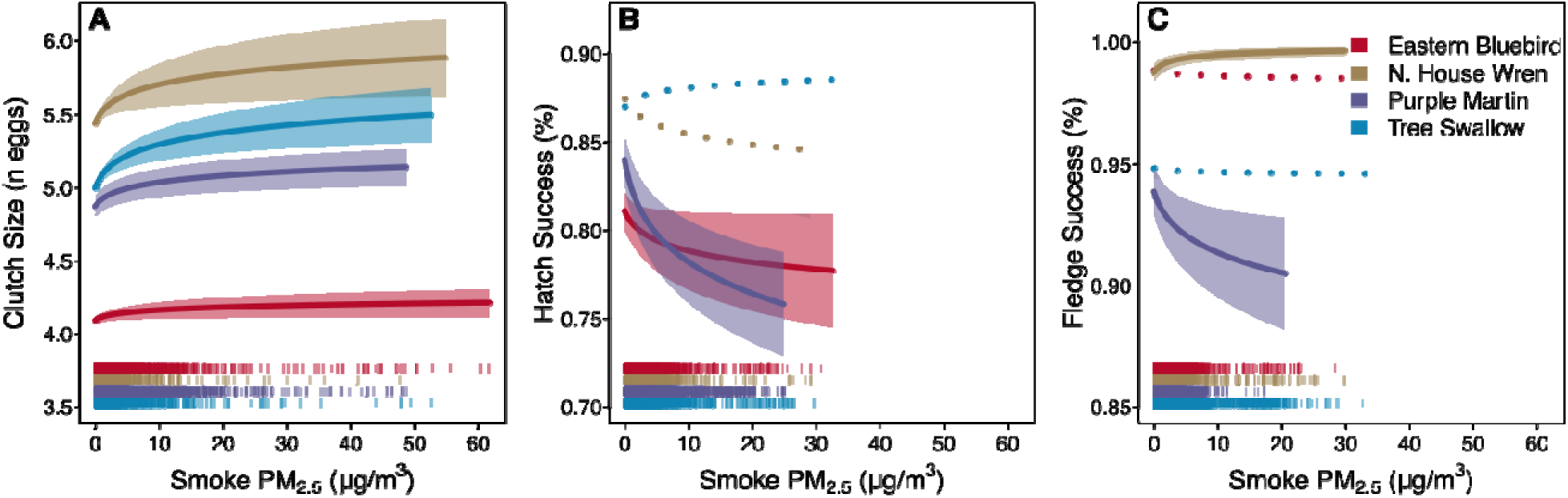
Smoke, controlling for site-specific spring climate, is associated with multiple fitness impacts to the avian reproductive cycle. Independent fitness responses include (**A**) clutch size, (**B**) hatch success, and (**C**) fledge success. Each response was tested independently across four species of cavity-nesting birds while controlling for maximum April temperature and total May precipitation at each site in each year, as well as the linear and quadratic phenological impact of day of year. Symbology as in fig. S4.

**Table S1.** Number of nest monitoring sites and temporal resampling of sites for each species after universal filtering. From the combined and filtered datasets, we limited nest monitoring sites within our study area to those with a minimum of three years of repeated data and a maximum of eight years within our study period (2018–2025).

| Species | Total sites | Temporal resampling (years; 2018-2025) of sites |  |  |  |  |  |
| --- | --- | --- | --- | --- | --- | --- | --- |
|  |  | 3 | 4 | 5 | 6 | 7 | 8 |
| Eastern Bluebird | 4115 | 1854 | 1150 | 553 | 338 | 167 | 53 |
| Northern House Wren | 1074 | 553 | 303 | 132 | 55 | 22 | 9 |
| Purple Martin | 1018 | 162 | 220 | 195 | 202 | 152 | 87 |
| Tree Swallow | 4294 | 1583 | 988 | 786 | 492 | 295 | 150 |

**Table S2.** Sample sizes of monitored nests per year used for analyses of each phenological or fitness response. Filtering steps and data requirements resulted in different numbers of monitored nests included per species per response per year, as well as a different total sample size of monitoring sites studied across all years. For all responses (excluding first lay date), repeat nesting attempts at the same site in a year were included.

| Species | Response | Number of nests monitored per year |  |  |  |  |  |  |  | Total number of sites |
| --- | --- | --- | --- | --- | --- | --- | --- | --- | --- | --- |
|  |  | 2018 | 2019 | 2020 | 2021 | 2022 | 2023 | 2024 | 2025 |  |
| Eastern Bluebird | First lay date | 1584 | 1858 | 1672 | 2280 | 2483 | 2568 | 2495 | 1608 | 4115 |
| Northern House Wren | First lay date | 449 | 510 | 546 | 615 | 546 | 557 | 470 | 394 | 1074 |
| Purple Martin | First lay date | 426 | 512 | 583 | 579 | 790 | 822 | 796 | 805 | 1018 |
| Tree Swallow | First lay date | 1830 | 2199 | 2157 | 2545 | 2583 | 2691 | 2504 | 2339 | 4294 |
| Eastern Bluebird | Incubation period | 1104 | 1264 | 1437 | 1586 | 1693 | 1746 | 1711 | 1352 | 2615 |
| Northern House Wren | Incubation period | 282 | 361 | 379 | 465 | 515 | 510 | 425 | 364 | 711 |
| Purple Martin | Incubation period | 365 | 425 | 477 | 458 | 610 | 618 | 641 | 620 | 845 |
| Tree Swallow | Incubation period | 1240 | 1524 | 1545 | 1878 | 2088 | 2088 | 1976 | 1851 | 3195 |
| Eastern Bluebird | Nestling period | 1220 | 1452 | 1353 | 1832 | 1910 | 2026 | 1963 | 1240 | 2890 |
| Northern House Wren | Nestling period | 241 | 316 | 329 | 408 | 471 | 462 | 385 | 310 | 645 |
| Purple Martin | Nestling period | 1496 | 1933 | 2309 | 2483 | 2953 | 3457 | 2921 | 3050 | 926 |
| Tree Swallow | Nestling period | 1126 | 1417 | 1462 | 1725 | 1900 | 1863 | 1778 | 1683 | 2969 |
| Eastern Bluebird | Clutch size | 1949 | 2258 | 2032 | 2787 | 2948 | 3078 | 3020 | 1874 | 4114 |
| Northern House Wren | Clutch size | 571 | 675 | 693 | 793 | 695 | 732 | 596 | 507 | 1073 |
| Purple Martin | Clutch size | 1739 | 2356 | 2627 | 2920 | 3650 | 4218 | 3609 | 4250 | 1018 |
| Tree Swallow | Clutch size | 1984 | 2345 | 2281 | 2776 | 2763 | 2903 | 2712 | 2623 | 4294 |
| Eastern Bluebird | Hatch success | 1867 | 2172 | 1922 | 2673 | 2841 | 2943 | 2891 | 1786 | 3948 |
| Northern House Wren | Hatch success | 520 | 636 | 648 | 736 | 647 | 673 | 551 | 470 | 1004 |
| Purple Martin | Hatch success | 1736 | 2351 | 2620 | 2917 | 3645 | 4213 | 3607 | 4244 | 1015 |
| Tree Swallow | Hatch success | 1926 | 2251 | 2189 | 2649 | 2620 | 2750 | 2511 | 2461 | 4112 |
| Eastern Bluebird | Fledge success | 1427 | 1641 | 1478 | 2068 | 2170 | 2301 | 2247 | 1361 | 3216 |
| Northern House Wren | Fledge success | 383 | 480 | 490 | 590 | 523 | 524 | 441 | 357 | 824 |
| Purple Martin | Fledge success | 1584 | 2073 | 2385 | 2593 | 3250 | 3699 | 3233 | 3384 | 963 |
| Tree Swallow | Fledge success | 1553 | 1837 | 1831 | 2152 | 2199 | 2283 | 2060 | 1939 | 3560 |

**Table S3.** Parameter results from generalized linear mixed models analyzing the association between pre-laying period smoke (PM_2.5_) and first lay date. Each species was modeled independently. Lay date was modeled as a gaussian-distributed random variable with an identity link and year and nest site as random effects. For each parameter, mean estimates are provided along with 95% confidence intervals and associated p-values.

| Species | Sample size | Parameter | Estimate | Lower 95% CI | Upper 95% CI | P-value |
| --- | --- | --- | --- | --- | --- | --- |
| Eastern Bluebird | 16548 | Intercept | 153.8 | 152.9 | 154.8 | < 0.001 |
| - | - | PM<sub>2.5</sub> | 2.458 | 2.032 | 2.884 | < 0.001 |
| Northern House Wren | 4087 | Intercept | 155.5 | 152.9 | 158.2 | < 0.001 |
| - | - | PM<sub>2.5</sub> | 1.669 | 0.663 | 2.676 | < 0.001 |
| Purple Martin | 5313 | Intercept | 148.1 | 146.0 | 150.3 | < 0.001 |
| - | - | PM<sub>2.5</sub> | 0.845 | 0.583 | 1.106 | < 0.001 |
| Tree Swallow | 18848 | Intercept | 137.1 | 135.6 | 138.6 | < 0.001 |
| - | - | PM<sub>2.5</sub> | 1.416 | 1.246 | 1.586 | < 0.001 |

**Table S4.**
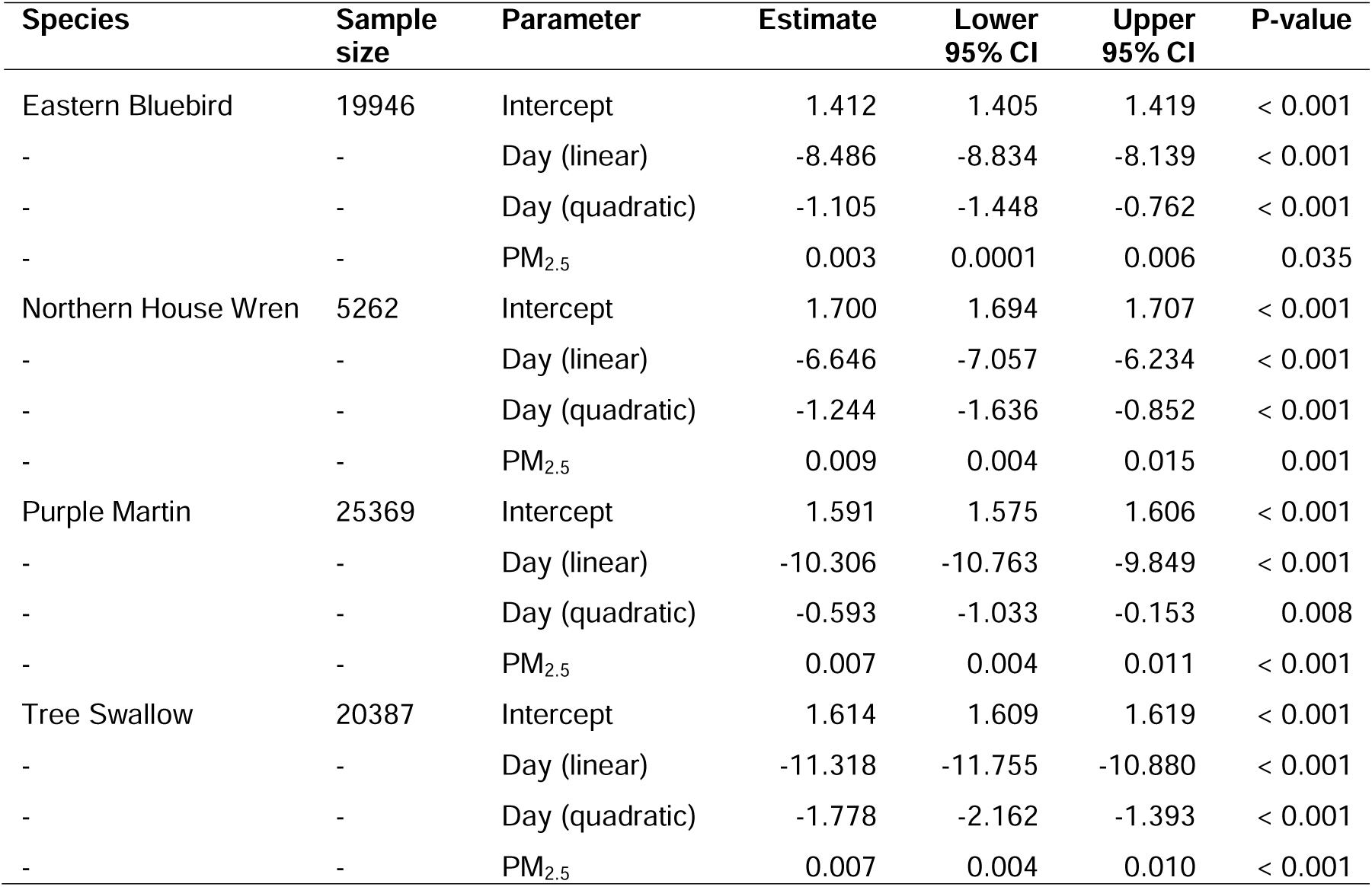
Parameter results from generalized linear mixed models analyzing the association between laying period smoke (PM_2.5_) and clutch size. Each species was modeled independently. Clutch size was modeled as a zero-truncated Poisson-distributed random variable with a log link and year and nest site as random effects. For each parameter, mean estimates are provided along with 95% confidence intervals and associated p-values.

**Table S5.** Parameter results from generalized linear mixed models analyzing the association between incubation period smoke (PM_2.5_) and incubation period duration. Each species was modeled independently. Incubation period was modeled as a zero-truncated Poisson-distributed random variable with a log link and year and nest site as random effects. For each parameter, mean estimates are provided along with 95% confidence intervals and associated p-values.

| Species | Sample size | Parameter | Estimate | Lower 95% CI | Upper 95% CI | P-value |
| --- | --- | --- | --- | --- | --- | --- |
| Eastern Bluebird | 11893 | Intercept | 2.532 | 2.528 | 2.536 | < 0.001 |
| - | - | Day (linear) | -1.631 | -1.962 | -1.299 | < 0.001 |
| - | - | Day (quadratic) | 0.053 | -0.279 | 0.385 | 0.755 |
| - | - | PM<sub>2.5</sub> | 0.005 | 0.002 | 0.008 | < 0.001 |
| Northern House Wren | 3301 | Intercept | 2.487 | 2.472 | 2.503 | < 0.001 |
| - | - | Day (linear) | -1.309 | -1.727 | -0.891 | < 0.001 |
| - | - | Day (quadratic) | 0.721 | 0.323 | 1.119 | < 0.001 |
| - | - | PM<sub>2.5</sub> | 0.012 | 0.002 | 0.021 | 0.017 |
| Purple Martin | 4214 | Intercept | 2.674 | 2.664 | 2.685 | < 0.001 |
| - | - | Day (linear) | -0.967 | -1.235 | -0.698 | < 0.001 |
| - | - | Day (quadratic) | 0.448 | 0.195 | 0.702 | 0.001 |
| - | - | PM<sub>2.5</sub> | 0.006 | -0.001 | 0.013 | 0.114 |
| Tree Swallow | 14190 | Intercept | 2.589 | 2.571 | 2.607 | < 0.001 |
| - | - | Day (linear) | -3.898 | -4.303 | -3.494 | < 0.001 |
| - | - | Day (quadratic) | 1.844 | 1.430 | 2.259 | < 0.001 |
| - | - | PM<sub>2.5</sub> | 0.028 | 0.024 | 0.032 | < 0.001 |

**Table S6.** Parameter results from generalized linear mixed models analyzing the association between incubation period smoke (PM_2.5_) and hatching success. Each species was modeled independently. Hatching success was modeled as a binomial-distributed random variable proportional to clutch size with a logit link and year and nest site as random effects. For each parameter, mean estimates are provided along with 95% confidence intervals and associated p-values.

| Species | Sample size | Parameter | Estimate | Lower 95% CI | Upper 95% CI | P-value |
| --- | --- | --- | --- | --- | --- | --- |
| Eastern Bluebird | 19095 | Intercept | 1.448 | 1.392 | 1.505 | < 0.001 |
| - | - | Day (linear) | -12.084 | -14.813 | -9.355 | < 0.001 |
| - | - | Day (quadratic) | -0.480 | -3.196 | 2.236 | 0.729 |
| - | - | PM<sub>2.5</sub> | -0.030 | -0.056 | -0.004 | 0.024 |
| Northern House Wren | 4881 | Intercept | 1.909 | 1.794 | 2.024 | < 0.001 |
| - | - | Day (linear) | 2.460 | -0.303 | 5.222 | 0.081 |
| - | - | Day (quadratic) | 0.525 | -2.118 | 3.167 | 0.697 |
| - | - | PM<sub>2.5</sub> | -0.041 | -0.087 | 0.004 | 0.077 |
| Purple Martin | 25333 | Intercept | 1.583 | 1.454 | 1.712 | < 0.001 |
| - | - | Day (linear) | -5.416 | -8.199 | -2.633 | < 0.001 |
| - | - | Day (quadratic) | -26.497 | -29.002 | -23.993 | < 0.001 |
| - | - | PM<sub>2.5</sub> | -0.100 | -0.127 | -0.072 | < 0.001 |
| Tree Swallow | 19357 | Intercept | 1.919 | 1.826 | 2.012 | < 0.001 |
| - | - | Day (linear) | -19.794 | -22.605 | -16.982 | < 0.001 |
| - | - | Day (quadratic) | -18.391 | -21.333 | -15.450 | < 0.001 |
| - | - | PM<sub>2.5</sub> | 0.014 | -0.010 | 0.039 | 0.258 |

**Table S7.** Parameter results from generalized linear mixed models analyzing the association between nestling period smoke (PM_2.5_) and nestling period duration. Each species was modeled independently. Nestling period was modeled as a zero-truncated Poisson-distributed random variable with a log link year and nest site as random effects. For each parameter, mean estimates are provided along with 95% confidence intervals and associated p-values.

| Species | Sample size | Parameter | Estimate | Lower 95% CI | Upper 95% CI | P-value |
| --- | --- | --- | --- | --- | --- | --- |
| Eastern Bluebird | 12996 | Intercept | 2.834 | 2.827 | 2.841 | < 0.001 |
| - | - | Day (linear) | -2.230 | -2.597 | -1.864 | < 0.001 |
| - | - | Day (quadratic) | 0.083 | -0.284 | 0.450 | 0.657 |
| - | - | PM<sub>2.5</sub> | 0.010 | 0.006 | 0.015 | < 0.001 |
| Northern House Wren | 2922 | Intercept | 2.744 | 2.735 | 2.752 | < 0.001 |
| - | - | Day (linear) | -0.942 | -1.292 | -0.591 | < 0.001 |
| - | - | Day (quadratic) | 0.010 | -0.340 | 0.361 | 0.955 |
| - | - | PM<sub>2.5</sub> | 0.006 | 0.000 | 0.013 | 0.064 |
| Purple Martin | 20602 | Intercept | 3.302 | 3.289 | 3.314 | < 0.001 |
| - | - | Day (linear) | -5.028 | -5.389 | -4.667 | < 0.001 |
| - | - | Day (quadratic) | -0.554 | -0.869 | -0.239 | 0.001 |
| - | - | PM<sub>2.5</sub> | 0.028 | 0.024 | 0.032 | < 0.001 |
| Tree Swallow | 12954 | Intercept | 2.954 | 2.946 | 2.962 | < 0.001 |
| - | - | Day (linear) | -2.403 | -2.703 | -2.103 | < 0.001 |
| - | - | Day (quadratic) | 1.477 | 1.167 | 1.788 | < 0.001 |
| - | - | PM<sub>2.5</sub> | -0.005 | -0.010 | -0.001 | 0.049 |

**Table S8.** Parameter results from generalized linear mixed models analyzing the association between nestling period smoke (PM_2.5_) and fledging success. Each species was modeled independently. Fledging success was modeled as a binomial-distributed random variable proportional to number of eggs hatched with a logit link year and nest site as random effects. For each parameter, mean estimates are provided along with 95% confidence intervals and associated p-values.

| Species | Sample size | Parameter | Estimate | Lower 95% CI | Upper 95% CI | P-value |
| --- | --- | --- | --- | --- | --- | --- |
| Eastern Bluebird | 14693 | Intercept | 4.446 | 4.278 | 4.615 | < 0.001 |
| - | - | Day (linear) | 10.615 | 5.239 | 15.990 | < 0.001 |
| - | - | Day (quadratic) | -5.040 | -10.323 | 0.243 | 0.062 |
| - | - | PM<sub>2.5</sub> | -0.017 | -0.059 | 0.026 | 0.440 |
| Northern House Wren | 3788 | Intercept | 4.600 | 4.260 | 4.939 | < 0.001 |
| - | - | Day (linear) | -2.710 | -7.296 | 1.877 | 0.247 |
| - | - | Day (quadratic) | -1.521 | -5.835 | 2.794 | 0.490 |
| - | - | PM<sub>2.5</sub> | 0.169 | 0.089 | 0.249 | < 0.001 |
| Purple Martin | 22201 | Intercept | 2.639 | 2.501 | 2.777 | < 0.001 |
| - | - | Day (linear) | -14.232 | -17.948 | -10.517 | < 0.001 |
| - | - | Day (quadratic) | 3.767 | 0.520 | 7.014 | 0.023 |
| - | - | PM<sub>2.5</sub> | -0.094 | -0.134 | -0.054 | < 0.001 |
| Tree Swallow | 15854 | Intercept | 2.905 | 2.822 | 2.987 | < 0.001 |
| - | - | Day (linear) | 13.507 | 9.486 | 17.527 | < 0.001 |
| - | - | Day (quadratic) | 4.556 | 0.166 | 8.946 | 0.042 |
| - | - | PM<sub>2.5</sub> | -0.030 | -0.073 | 0.013 | 0.178 |

**Table S9.**
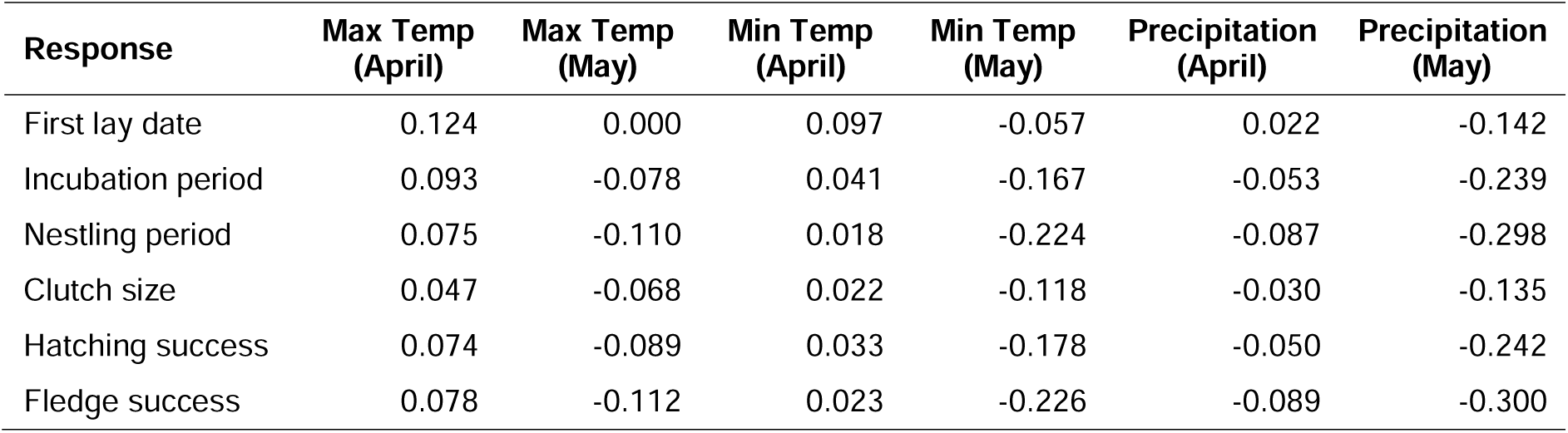
Correlations between climatic variables and smoke exposure for each reproductive response. Pearson correlations were calculated combining nest data from all species across all years (2018–2025).

**Table S10.** Parameter results from generalized linear mixed models analyzing the association between pre-laying period smoke (PM_2.5_) and first lay date, controlling for spring climate. Each species was modeled independently. Lay date was modeled as a gaussian-distributed random variable with an identity link and year and nest site as random effects. For each parameter, mean estimates are provided along with 95% confidence intervals and associated p-values.

| Species | Sample size | Parameter | Estimate | Lower 95% CI | Upper 95% CI | P-value |
| --- | --- | --- | --- | --- | --- | --- |
| Eastern Bluebird | 16548 | Intercept | 153.8 | 152.8 | 154.7 | < 0.001 |
| - | - | PM<sub>2.5</sub> | 2.683 | 2.257 | 3.109 | < 0.001 |
| - | - | Tmax (Apr) | -1.614 | -1.950 | -1.279 | < 0.001 |
| - | - | Precip (May) | 0.366 | -0.002 | 0.734 | 0.051 |
| Northern House Wren | 4087 | Intercept | 155.5 | 153.7 | 157.3 | < 0.001 |
| - | - | PM<sub>2.5</sub> | 1.851 | 0.856 | 2.846 | < 0.001 |
| - | - | Tmax (Apr) | -3.669 | -4.703 | -2.634 | < 0.001 |
| - | - | Precip (May) | -0.293 | -1.192 | 0.606 | 0.523 |
| Purple Martin | 5313 | Intercept | 147.8 | 145.0 | 150.7 | < 0.001 |
| - | - | PM<sub>2.5</sub> | 1.334 | 1.078 | 1.591 | < 0.001 |
| - | - | Tmax (Apr) | -6.009 | -6.406 | -5.613 | < 0.001 |
| - | - | Precip (May) | 0.544 | 0.175 | 0.913 | 0.004 |
| Tree Swallow | 18848 | Intercept | 137.0 | 135.5 | 138.5 | < 0.001 |
| - | - | PM<sub>2.5</sub> | 1.502 | 1.332 | 1.673 | < 0.001 |
| - | - | Tmax (Apr) | -1.284 | -1.492 | -1.076 | < 0.001 |
| - | - | Precip (May) | 0.233 | 0.008 | 0.457 | 0.042 |

**Table S11.** Parameter results from generalized linear mixed models analyzing the association between incubation period smoke (PM_2.5_) and incubation period duration, controlling for spring climate. Each species was modeled independently. Incubation period was modeled as a zero-truncated Poisson-distributed random variable with a log link and year and nest site as random effects. For each parameter, mean estimates are provided along with 95% confidence intervals and associated p-values.

| Species | Sample size | Parameter | Estimate | Lower 95% CI | Upper 95% CI | P-value |
| --- | --- | --- | --- | --- | --- | --- |
| Eastern Bluebird | 11893 | Intercept | 2.532 | 2.528 | 2.536 | < 0.001 |
| - | - | PM<sub>2.5</sub> | 0.005 | 0.002 | 0.008 | 0.001 |
| - | - | Tmax (Apr) | 0.001 | -0.002 | 0.005 | 0.534 |
| - | - | Precip (May) | -0.001 | -0.004 | 0.003 | 0.704 |
| - | - | Day (linear) | -1.622 | -1.955 | -1.290 | < 0.001 |
| - | - | Day (quadratic) | 0.059 | -0.274 | 0.391 | 0.729 |
| Northern House Wren | 3301 | Intercept | 2.486 | 2.474 | 2.497 | < 0.001 |
| - | - | PM<sub>2.5</sub> | 0.013 | 0.004 | 0.022 | 0.003 |
| - | - | Tmax (Apr) | -0.022 | -0.031 | -0.013 | < 0.001 |
| - | - | Precip (May) | 0.008 | -0.001 | 0.017 | 0.069 |
| - | - | Day (linear) | -1.371 | -1.786 | -0.955 | < 0.001 |
| - | - | Day (quadratic) | 0.772 | 0.374 | 1.169 | < 0.001 |
| Purple Martin | 4214 | Intercept | 2.674 | 2.664 | 2.684 | < 0.001 |
| - | - | PM<sub>2.5</sub> | 0.008 | 0.001 | 0.015 | 0.031 |
| - | - | Tmax (Apr) | -0.001 | -0.008 | 0.005 | 0.656 |
| - | - | Precip (May) | 0.008 | 0.002 | 0.015 | 0.010 |
| - | - | Day (linear) | -0.960 | -1.258 | -0.662 | < 0.001 |
| - | - | Day (quadratic) | 0.437 | 0.175 | 0.699 | 0.001 |
| Tree Swallow | 14190 | Intercept | 2.589 | 2.572 | 2.606 | < 0.001 |
| - | - | PM<sub>2.5</sub> | 0.029 | 0.024 | 0.033 | < 0.001 |
| - | - | Tmax (Apr) | 0.000 | -0.005 | 0.005 | 0.942 |
| - | - | Precip (May) | 0.004 | 0.000 | 0.009 | 0.078 |
| - | - | Day (linear) | -3.896 | -4.301 | -3.490 | < 0.001 |
| - | - | Day (quadratic) | 1.823 | 1.398 | 2.248 | < 0.001 |

**Table S12.** Parameter results from generalized linear mixed models analyzing the association between nestling period smoke (PM_2.5_) and nestling period duration, controlling for spring climate. Each species was modeled independently. Nestling period was modeled as a zero-truncated Poisson-distributed random variable with a log link and year and nest site as random effects. For each parameter, mean estimates are provided along with 95% confidence intervals and associated p-values.

| Species | Sample size | Parameter | Estimate | Lower 95% CI | Upper 95% CI | P-value |
| --- | --- | --- | --- | --- | --- | --- |
| Eastern Bluebird | 12996 | Intercept | 2.833 | 2.827 | 2.839 | < 0.001 |
| - | - | PM<sub>2.5</sub> | 0.009 | 0.005 | 0.013 | < 0.001 |
| - | - | Tmax (Apr) | -0.024 | -0.028 | -0.020 | < 0.001 |
| - | - | Precip (May) | 0.003 | -0.001 | 0.007 | 0.137 |
| - | - | Day (linear) | -2.333 | -2.699 | -1.966 | < 0.001 |
| - | - | Day (quadratic) | -0.029 | -0.396 | 0.339 | 0.879 |
| Northern House Wren | 2922 | Intercept | 2.743 | 2.735 | 2.751 | < 0.001 |
| - | - | PM<sub>2.5</sub> | 0.009 | 0.002 | 0.017 | 0.015 |
| - | - | Tmax (Apr) | -0.014 | -0.022 | -0.006 | < 0.001 |
| - | - | Precip (May) | -0.004 | -0.011 | 0.003 | 0.290 |
| - | - | Day (linear) | -1.002 | -1.352 | -0.652 | < 0.001 |
| - | - | Day (quadratic) | 0.066 | -0.284 | 0.417 | 0.710 |
| Purple Martin | 20602 | Intercept | 3.300 | 3.286 | 3.314 | < 0.001 |
| - | - | PM<sub>2.5</sub> | 0.028 | 0.024 | 0.032 | < 0.001 |
| - | - | Tmax (Apr) | -0.025 | -0.030 | -0.019 | < 0.001 |
| - | - | Precip (May) | 0.004 | 0.001 | 0.007 | 0.011 |
| - | - | Day (linear) | -5.432 | -5.801 | -5.064 | < 0.001 |
| - | - | Day (quadratic) | -0.342 | -0.659 | -0.026 | 0.034 |
| Tree Swallow | 12954 | Intercept | 2.953 | 2.944 | 2.963 | < 0.001 |
| - | - | PM<sub>2.5</sub> | 0.000 | -0.006 | 0.005 | 0.940 |
| - | - | Tmax (Apr) | -0.012 | -0.016 | -0.009 | < 0.001 |
| - | - | Precip (May) | 0.010 | 0.006 | 0.014 | < 0.001 |
| - | - | Day (linear) | -2.418 | -2.718 | -2.118 | < 0.001 |
| - | - | Day (quadratic) | 1.638 | 1.322 | 1.955 | < 0.001 |

**Table S13.**
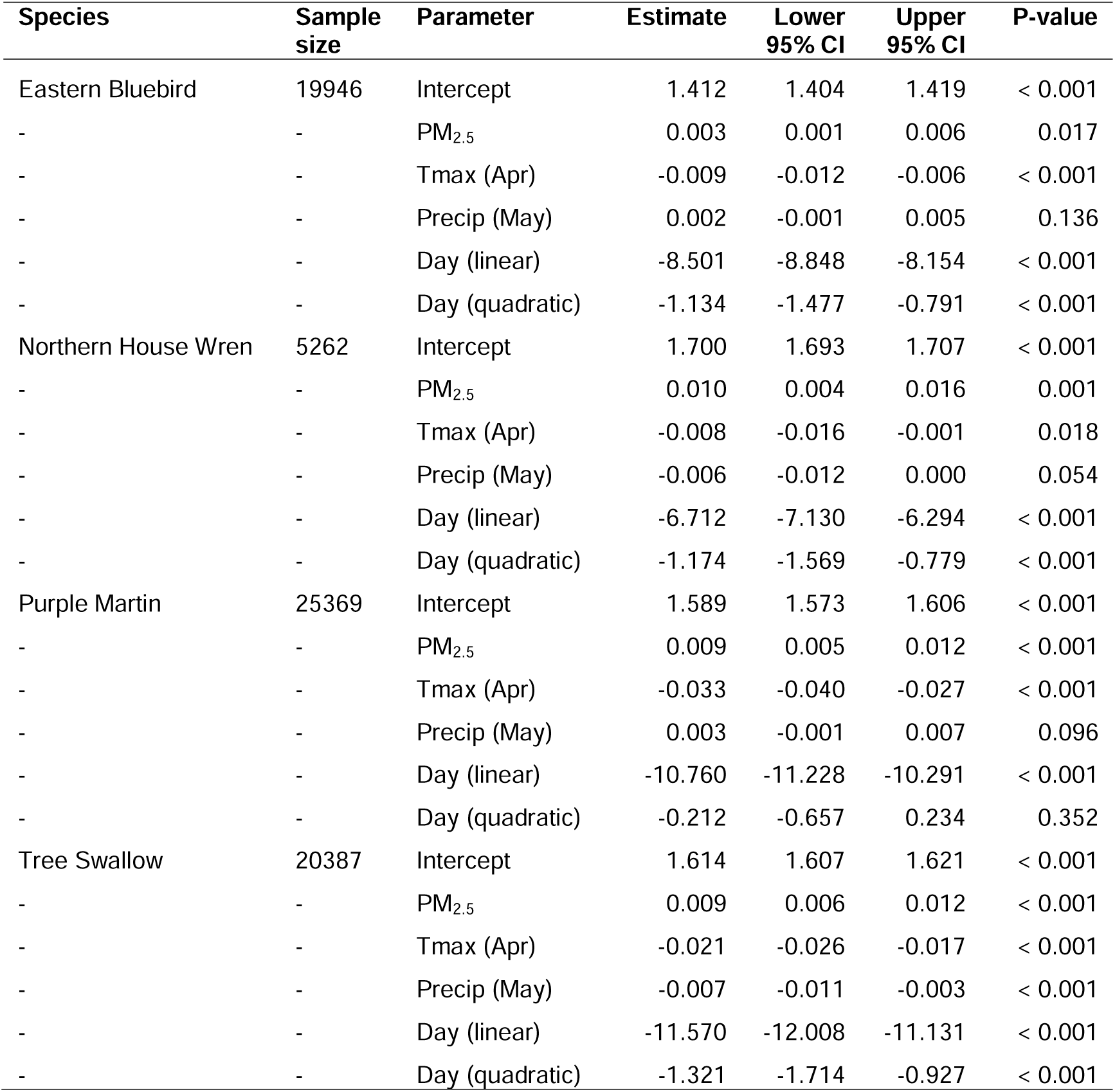
Parameter results from generalized linear mixed models analyzing the association between laying period smoke (PM_2.5_) and clutch size, controlling for spring climate. Each species was modeled independently. Clutch size was modeled as a zero-truncated Poisson-distributed random variable with log link and year and nest site as random effects. For each parameter, mean estimates are provided along with 95% confidence intervals and associated p-values.

**Table S14.** Parameter results from generalized linear mixed models analyzing the association between incubation period smoke (PM_2.5_) and hatching success, controlling for spring climate. Each species was modeled independently. Hatching success was modeled as a binomial-distributed random variable proportional to clutch size with a logit link and year and nest site as random effects. For each parameter, mean estimates are provided along with 95% confidence intervals and associated p-values.

| Species | Sample size | Parameter | Estimate | Lower 95% CI | Upper 95% CI | P-value |
| --- | --- | --- | --- | --- | --- | --- |
| Eastern Bluebird | 19095 | Intercept | 1.434 | 1.359 | 1.508 | < 0.001 |
| - | - | PM<sub>2.5</sub> | -0.029 | -0.056 | -0.002 | 0.033 |
| - | - | Tmax (Apr) | -0.242 | -0.281 | -0.203 | < 0.001 |
| - | - | Precip (May) | 0.018 | -0.006 | 0.043 | 0.145 |
| - | - | Day (linear) | -12.279 | -15.007 | -9.551 | < 0.001 |
| - | - | Day (quadratic) | -1.094 | -3.810 | 1.622 | 0.430 |
| Northern House Wren | 4881 | Intercept | 1.912 | 1.763 | 2.061 | < 0.001 |
| - | - | PM<sub>2.5</sub> | -0.040 | -0.088 | 0.008 | 0.106 |
| - | - | Tmax (Apr) | -0.148 | -0.242 | -0.054 | 0.002 |
| - | - | Precip (May) | -0.100 | -0.155 | -0.046 | < 0.001 |
| - | - | Day (linear) | 2.265 | -0.517 | 5.046 | 0.111 |
| - | - | Day (quadratic) | 0.707 | -1.944 | 3.357 | 0.601 |
| Purple Martin | 25333 | Intercept | 1.580 | 1.468 | 1.692 | < 0.001 |
| - | - | PM<sub>2.5</sub> | -0.108 | -0.135 | -0.080 | < 0.001 |
| - | - | Tmax (Apr) | -0.012 | -0.065 | 0.040 | 0.645 |
| - | - | Precip (May) | -0.107 | -0.130 | -0.083 | < 0.001 |
| - | - | Day (linear) | -5.352 | -8.172 | -2.532 | < 0.001 |
| - | - | Day (quadratic) | -26.745 | -29.274 | -24.216 | < 0.001 |
| Tree Swallow | 19357 | Intercept | 1.916 | 1.814 | 2.018 | < 0.001 |
| - | - | PM<sub>2.5</sub> | 0.019 | -0.006 | 0.044 | 0.136 |
| - | - | Tmax (Apr) | -0.053 | -0.095 | -0.012 | 0.012 |
| - | - | Precip (May) | 0.033 | 0.003 | 0.062 | 0.029 |
| - | - | Day (linear) | -20.133 | -22.953 | -17.313 | < 0.001 |
| - | - | Day (quadratic) | -17.879 | -20.846 | -14.911 | < 0.001 |

**Table S15.** Parameter results from generalized linear mixed models analyzing the association between nestling period smoke (PM_2.5_) and fledging success, controlling for spring climate. Each species was modeled independently. Fledging success was modeled as a binomial-distributed random variable proportional to number of eggs hatched with a logit link and year and nest site as random effects. For each parameter, mean estimates are provided along with 95% confidence intervals and associated p-values.

| Species | Sample size | Parameter | Estimate | Lower 95% CI | Upper 95% CI | P-value |
| --- | --- | --- | --- | --- | --- | --- |
| Eastern Bluebird | 14693 | Intercept | 4.439 | 4.272 | 4.605 | < 0.001 |
| - | - | PM<sub>2.5</sub> | -0.036 | -0.080 | 0.008 | 0.105 |
| - | - | Tmax (Apr) | 0.180 | 0.094 | 0.266 | < 0.001 |
| - | - | Precip (May) | -0.010 | -0.057 | 0.037 | 0.675 |
| - | - | Day (linear) | 10.874 | 5.500 | 16.248 | < 0.001 |
| - | - | Day (quadratic) | -4.429 | -9.741 | 0.884 | 0.102 |
| Northern House Wren | 3788 | Intercept | 4.583 | 4.249 | 4.917 | < 0.001 |
| - | - | PM<sub>2.5</sub> | 0.233 | 0.146 | 0.319 | < 0.001 |
| - | - | Tmax (Apr) | -0.160 | -0.277 | -0.043 | 0.007 |
| - | - | Precip (May) | 0.066 | -0.010 | 0.141 | 0.089 |
| - | - | Day (linear) | -3.229 | -7.833 | 1.375 | 0.169 |
| - | - | Day (quadratic) | -1.299 | -5.622 | 3.025 | 0.556 |
| Purple Martin | 22201 | Intercept | 2.648 | 2.470 | 2.827 | < 0.001 |
| - | - | PM<sub>2.5</sub> | -0.082 | -0.122 | -0.041 | < 0.001 |
| - | - | Tmax (Apr) | -0.415 | -0.492 | -0.338 | < 0.001 |
| - | - | Precip (May) | -0.125 | -0.158 | -0.091 | < 0.001 |
| - | - | Day (linear) | -16.900 | -20.638 | -13.163 | < 0.001 |
| - | - | Day (quadratic) | 5.839 | 2.546 | 9.132 | 0.001 |
| Tree Swallow | 15854 | Intercept | 2.903 | 2.819 | 2.988 | < 0.001 |
| - | - | PM<sub>2.5</sub> | -0.008 | -0.057 | 0.040 | 0.731 |
| - | - | Tmax (Apr) | -0.141 | -0.198 | -0.084 | < 0.001 |
| - | - | Precip (May) | -0.051 | -0.091 | -0.011 | 0.012 |
| - | - | Day (linear) | 13.285 | 9.257 | 17.313 | < 0.001 |
| - | - | Day (quadratic) | 5.955 | 1.508 | 10.402 | 0.009 |

